# Dynamic Responses to an Inflammatory Challenge Distinguish Metabolic Health Across Lean and Obese Individuals

**DOI:** 10.64898/2026.09.15.751578

**Authors:** Saeideh Tavajoh, Bianca E. Suur, Madison Clark, Adriana A. Becerril-Campos, Elia Velasco, Brigita Medelytė, Line Boel Nørregaard, Stine Julie Tingskov, Amanda Bæk, Lykke Skaarup, Erik Schrøder, Maryam Dost, Sigrid Bjerge Gribsholt, Jens M. Bruun, Marianne Quiding Järbrink, Sofia Nyström, Ida Bergström, Catherine Åhlund, Jane Palsgaard Pedersen, Niels Jessen, Lin Lin, Andras Harazin, Henrik H. Thomsen, Per-Anders Jansson, Robert Blomgran, Stephan Lange, Matúš Soták, Emma Börgeson

**Author notes:** Corresponding author; Emma Börgeson, Aarhus University, The Skou Building, Høegh-Guldbergs Gade 10, DK-8000 Aarhus C, Denmark. Co-first authorship (Saeideh Tavajoh, Bianca E. Suur). Co-senior authorship (Matúš Soták, Emma Börgeson).

## Abstract

Inflammation is a key driver of cardiometabolic disease, yet it remains unclear whether systemic inflammatory markers can distinguish metabolically healthy from unhealthy individuals or capture the temporal dynamics of inflammation. Here, we combined systemic immune profiling with a cantharidin-induced peripheral blister model to investigate dynamic inflammatory regulation across metabolic phenotypes in metabolically healthy lean (MHL), metabolically unhealthy lean (MUL), metabolically healthy obese (MHO), and metabolically unhealthy obese (MUO) individuals. While systemic inflammation was elevated in obesity, differences between metabolically healthy and unhealthy groups were modest, with limited discrimination by plasma proteomics, circulating leukocyte phenotyping, and whole-blood transcriptomics. In contrast, the dynamic response to inflammatory challenge revealed pronounced differences at proteomic, cellular, and transcriptomic levels. Metabolically unhealthy individuals exhibited exaggerated early innate immune responses, impaired inflammatory resolution and tissue repair, reduced recruitment of reparative immune cells, and sustained T cell presence. Transcriptomic analyses further showed blunted dynamic gene regulation and defective epidermal barrier restoration. These findings indicate that metabolic health is better reflected in tissue-level inflammatory dynamics than in systemic measures.

## Introduction

Obesity is a heterogeneous condition, with some individuals remaining metabolically healthy despite excess adiposity. This observation has led to stratification into distinct phenogroups of metabolically healthy obese (MHO) and metabolically unhealthy obese (MUO) (Blüher, 2020; Iacobini et al., 2019; Petersen et al., 2024). This concept also extends to lean individuals, who can be classified as metabolically healthy lean (MHL) or metabolically unhealthy lean (MUL) (Stefan et al., 2017; Zembic et al., 2021). Thus, adiposity per se is not the sole determinant of cardiometabolic risk, and MUL individuals show mortality outcomes comparable to MUO, whereas MHO individuals exhibit partial protection (Chen et al., 2024).

Low-grade inflammation is a defining feature of obesity and contributes to the increased risk of metabolic and cardiovascular complications, including diabetes and cardiovascular disease (Blüher, 2025; Emanuela et al., 2012; Soták et al., 2025). Inflammatory pathways have therefore become attractive therapeutic targets, with pharmacological inhibition of pro-inflammatory cytokines providing partial protection against cardiometabolic disease (Dominguez et al., 2005; Li et al., 2023; Ridker et al., 2021; Ridker et al., 2017; Ridker et al., 2012; Stanley et al., 2011). However, human studies indicate that therapeutic responses are influenced by underlying inflammatory heterogeneity (Ridker et al., 2018; Soták et al., 2022). This suggests that obesity-associated inflammation is not uniform across individuals, and that patient stratification may be important for identifying those most likely to benefit from inflammation-targeted therapies.

Despite the established link between obesity and low-grade inflammation, it remains unclear whether inflammation differs between metabolically healthy and unhealthy phenogroups (Iglesias Molli et al., 2017; Spoto et al., 2023). Importantly, previous work has largely assessed inflammation using circulating markers, such as C-reactive protein (CRP), or tissue biopsies, which provide valuable but static measures of inflammatory status (Eswar et al., 2024; Petersen et al., 2024; Qu et al., 2014; Su et al., 2024). These approaches do not capture how inflammation is temporally regulated after an acute challenge; from initiation and amplification to active resolution. Because defective resolution can sustain inflammatory responses and promote chronic disease (Furman et al., 2019; Sugimoto et al., 2016), inflammatory kinetics may therefore reveal disease-relevant differences that are not apparent from baseline systemic markers alone. Indeed, emerging evidence suggests that impaired resolution underlies multiple pathological conditions (Fishbein et al., 2021; Fredman and MacNamara, 2021), and specialized pro-resolving mediators may hold therapeutic potential (Soták et al., 2025). Thus, temporal assessment of inflammatory responses in humans remains limited, and the dynamic regulation of inflammation across metabolic phenotypes is largely unexplored.

Here, we use a human blister-wound model to assess acute and resolving inflammation after a controlled insult across population-based cohorts stratified into distinct cardiometabolic phenotypes. By integrating the dynamic readout of the blister model with circulating immune profiling, plasma proteomics and transcriptomics, we compare local inflammatory kinetics with baseline systemic inflammatory status. Our findings reveal that the regulation of inflammation in a dynamic manner offers a more accurate reflection of metabolic health than systemic markers alone. This finding emphasizes the need to consider inflammatory kinetics to better understand the diverse inflammatory responses observed in patients.

## Results and Discussion

### Clinical characterization of distinct metabolic phenotypes

Volunteers were recruited from two national Scandinavian population-based cohort studies (**Fig. S1A**) and underwent detailed anthropometric and clinical characterization (**Table 1**). Based on BMI and prespecified metabolic criteria, participants were classified into four phenogroups: metabolically healthy lean (MHL), metabolically unhealthy lean (MUL), metabolically healthy obese (MHO) and metabolically unhealthy obese (MUO) (see **Materials and Methods**). The contribution of each criterion to phenogroup classification is shown in **Fig. S1B**. Notably, MHO individuals showed little evidence of hypertriglyceridemia, whereas hypertension was present across all phenogroups, including among MHL individuals (**Fig. S1B**). In contrast, the metabolically unhealthy phenogroups, MUL and MUO, were primarily characterized by low HDL cholesterol and hyperglycemia, as well as hypertriglyceridemia. MUO individuals had greater central adiposity and insulin resistance than MHO, and MUL individuals showed higher central adiposity than MHL despite a lean BMI, consistent with a “thin-on-the-outside, fat-on-the-inside” phenotype and indicating that adverse fat distribution is linked to metabolic risk across BMI categories (Salmón-Gómez et al., 2023; Thomas et al., 2012). Collectively, the phenogroups differed primarily in the expected cardiometabolic features used to define metabolic health, while other clinical characteristics were broadly comparable within each adiposity group (**Table 1),** consistent with previous observations (Anand et al., 2025; Spoto et al., 2023). Importantly, the groups were comparable in age and sex distribution (**Table 1**).

**Table 1:**
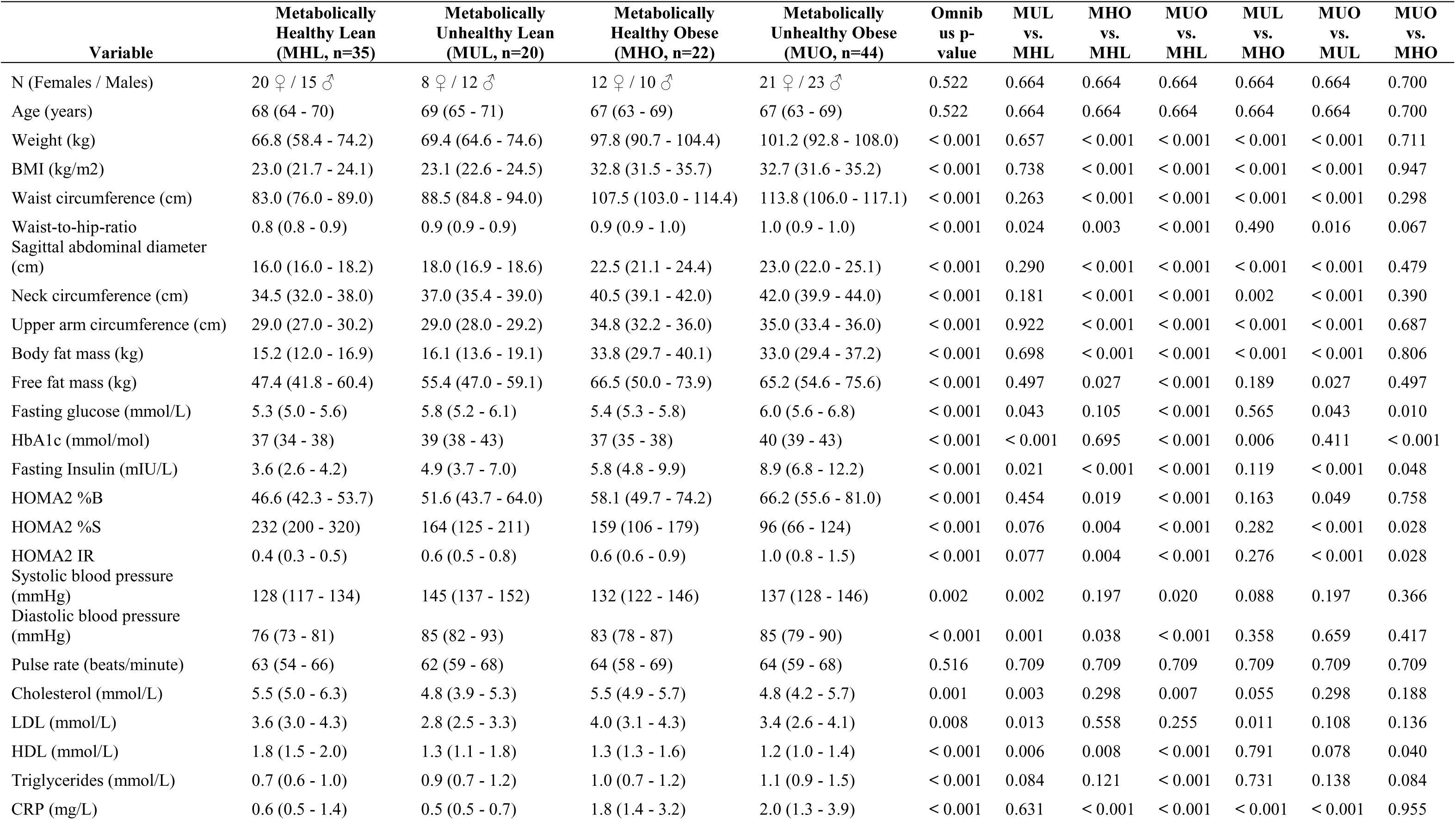
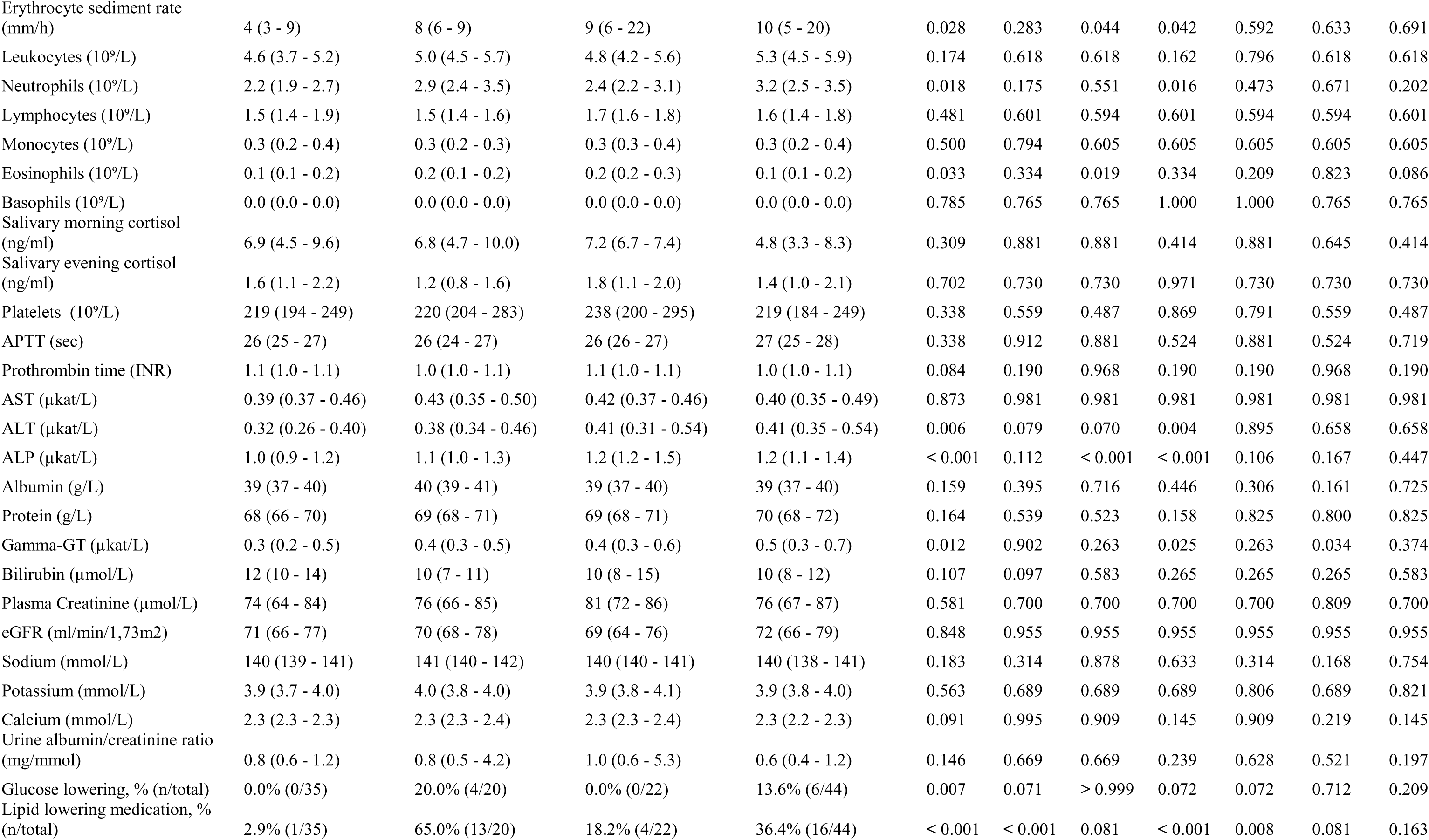
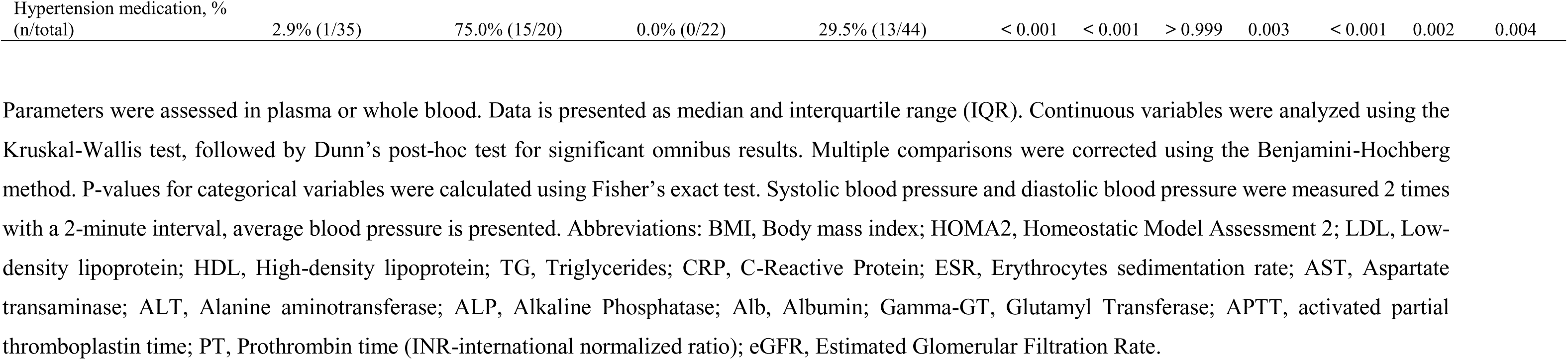
Clinical Characteristics of Participants. Parameters were assessed in plasma or whole blood. Data is presented as median and interquartile range (IQR). Continuous variables were analyzed using the Kruskal-Wallis test, followed by Dunn’s post-hoc test for significant omnibus results. Multiple comparisons were corrected using the Benjamini-Hochberg method. P-values for categorical variables were calculated using Fisher’s exact test. Systolic blood pressure and diastolic blood pressure were measured 2 times with a 2-minute interval, average blood pressure is presented. Abbreviations: BMI, Body mass index; HOMA2, Homeostatic Model Assessment 2; LDL, Low-density lipoprotein; HDL, High-density lipoprotein; TG, Triglycerides; CRP, C-Reactive Protein; ESR, Erythrocytes sedimentation rate; AST, Aspartate transaminase; ALT, Alanine aminotransferase; ALP, Alkaline Phosphatase; Alb, Albumin; Gamma-GT, Glutamyl Transferase; APTT, activated partial thromboplastin time; PT, Prothrombin time (INR-international normalized ratio); eGFR, Estimated Glomerular Filtration Rate.

### Metabolic dysfunction associates with larger blisters and altered vascular dynamics

To assess inflammatory regulation across lean and obese metabolic phenotypes, we used a cantharidin-induced blister-wound model that captures the response to a controlled inflammatory insult *in vivo* (De Maeyer et al., 2020; Jenner and Gilroy, 2012). Blister exudates were collected during both the acute inflammation phase (24 h post-initiation, immediately following the cantharidin challenge), and the resolution phase (72 h post-initiation, representing the 24-h challenge followed by 48 h of recovery). This serial sampling enabled time-resolved profiling of soluble mediators and infiltrating immune cells by multiplex protein assays, flow cytometry, and RNA sequencing, providing a multidimensional assessment of inflammatory dynamics (**Fig. 1A**).

**Figure 1.**
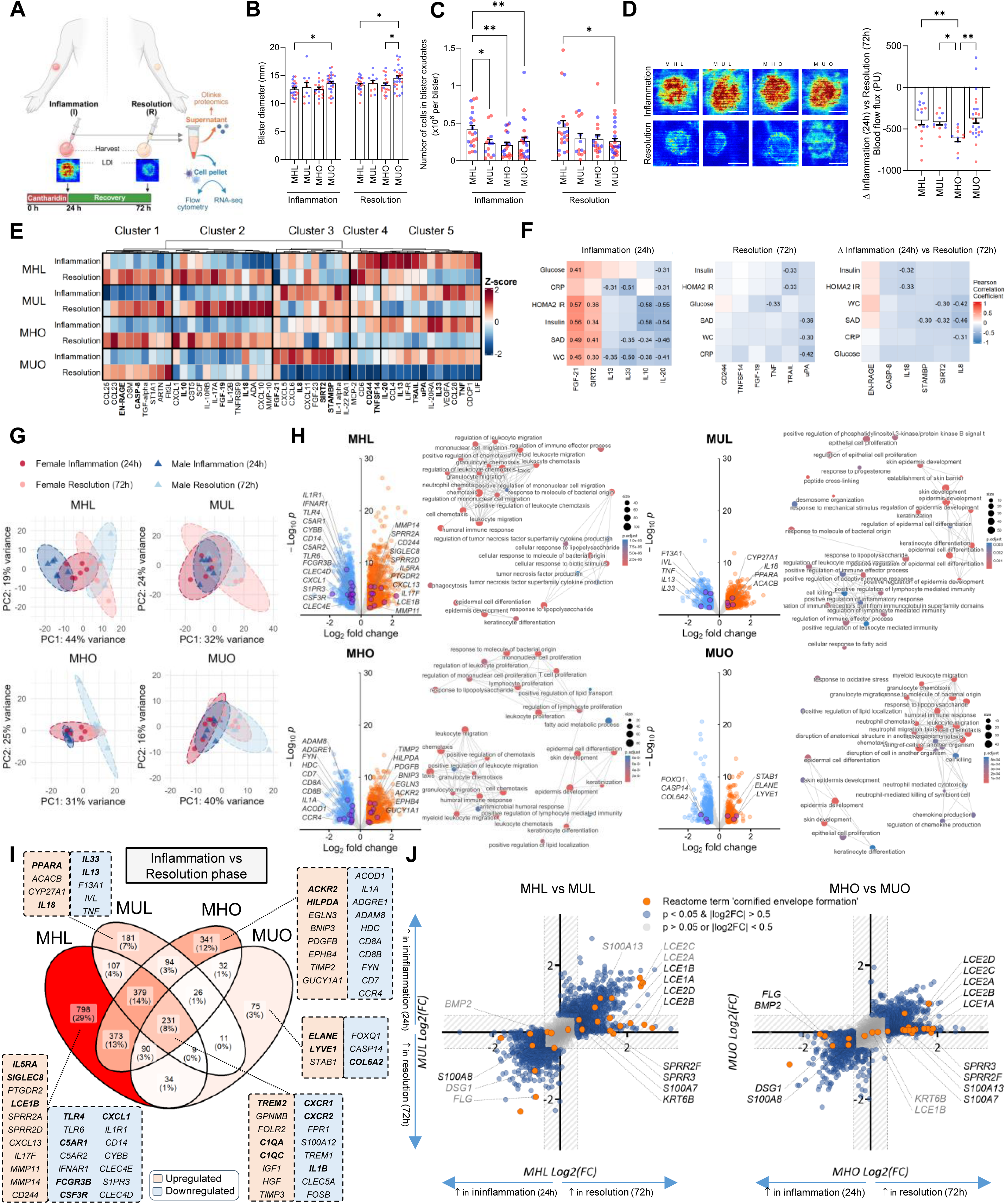
Cantharidin-induced blister profiling reveals metabolic phenotype–specific inflammatory dynamics. **(A)** Schematic overview of the cantharidin-induced skin blister model used to assess peripheral tissue inflammation during the acute inflammation phase, collected 24 h post-initiation immediately after inflammatory stimulation, and the resolution phase, collected 72 h post-initiation after inflammatory stimulation followed by a recovery period. Blister exudates were centrifuged, and fluid and cellular fractions were analyzed separately by Olink proteomics, flow cytometry, and RNA sequencing. **(B)** Blister size estimated by diameter measurement and shown in millimeters. **(C)** Number of live cells collected per blister. **(D)** Representative laser Doppler imaging scans at baseline and at 24 h and 72 h after cantharidin-induced blister formation. Mean flux, expressed as perfusion units (PU), was quantified from selected regions of interest using moorLDI Review v6.2.1, and the difference between the resolution and acute inflammation phases is shown. (**E**) Heatmap and hierarchical clustering of blister exudate proteins that either differed between metabolic phenotypes, inflammation/resolution phases, or ranked among the top contributors to group separation in PLS-DA analyses. Data are presented as column-wise scaled and centered means (Z-score) of NPX (normalized protein expression) values in log2 scale. Protein levels were modeled via LMMs (adjusted for age and sex; REML) with individuals as random effects. (**F**) Pearson correlation coefficients (*r*) between selected blister exudate proteins (with top VIP scores from PLS-DA analysis) and log2-transformed clinical markers of systemic inflammation (CRP), glycemic control (fasting glucose, insulin, HOMA2-IR), and adiposity (sagittal height, waist circumference). Results are illustrated as heatmaps displaying correlation for the acute inflammatory phase (24 h, left), the resolution phase (72 h; middle), and the longitudinal change (Δ log2 fold change, 72 h vs. 24 h; right). The color scale indicates the strength and direction of the correlation (red = positive, blue = negative). Correlation coefficients are numerically displayed only for statistically significant associations (p ≤ 0.05). **(G)** Principal component (PCA) analysis of blister cell-derived transcriptome across metabolic phenotypes based on the top 500 genes with the highest inter-sample variance. Axis labels indicate the percentage of total variance explained by each component. **(H)** Differential gene expression. Volcano plots illustrate global transcriptional changes across the four metabolic phenotypes. The x-axis represents stabilized log2 fold changes (LFC) and the y-axis denotes log10 adjusted p values. Dashed lines indicate significance thresholds (|LFC| > 0.5, adjusted p < 0.05). Regulated genes are shown in blue. The top 10 enriched Gene Ontology Biological Process (GO:BP) terms for each phenotype are shown, ranked by significance. Only protein-coding genes were included in the analysis. **(I)** Venn diagram illustrating the distribution of unique and shared differentially expressed genes (adjusted p < 0.05, |LFC| > 0.5) among the four metabolic phenotypes. Representative genes of interest are highlighted in colored boxes, where light red and light blue backgrounds denote up- and down-regulated transcripts, respectively. **(J)** Four-way plot contrasting stabilized log2 fold changes (LFC) in MHL vs. MUL (x-axis) and MHO vs. MUO (y-axis). Each point represents a single gene, with dashed lines indicating the significance thresholds (|LFC| = 0.5, adjusted p < 0.05). Positive values indicating higher expression during resolution and negative values indicating higher expression during acute inflammation. Genes positioned in the upper-right or lower-left quadrants showed concordant regulation between phenotypes, whereas genes in the opposing quadrants would have shown divergent phase-dependent regulation. Genes for top enriched Reactome term ‘cornified envelope formation’ are highlighted in orange. MHL, metabolically healthy lean; MUL, metabolically unhealthy lean; MHO, metabolically healthy obese; MUO, metabolically unhealthy obese. Data are presented as mean ± SEM. Statistical significance in B–D was calculated using Kruskal-Wallis test with Dunn’s post hoc test for pairwise comparisons. *p < 0.05, **p < 0.01, and ***p < 0.001.

We first assessed blister formation as a macroscopic readout of the inflammatory response. Although cantharidin-coated discs were uniform in size, MUO individuals developed larger blisters than the other groups (**Fig. 1B**). We next quantified cellular infiltration into the blister exudate (**Fig. 1C**). During the acute inflammation phase, all groups (MUL, MHO, and MUO) exhibited reduced numbers of infiltrating cells compared with MHL. In the resolution phase, these differences were less pronounced but remained significant in MUO relative to MHL (**Fig. 1C**).

Given that obesity and metabolic dysfunction have been linked to altered vascular and tissue responses (Mori et al., 2017), albeit with inconsistent associations with skin blood flow (Andreieva et al., 2021; Francischetti et al., 2011), we next measured perfusion at the blister site using laser Doppler imaging **(Fig. 1D**). Perfusion generally increased during acute inflammation and declined during resolution. However, this phase-to-phase difference was most pronounced in MHO individuals.

In summary, MUO individuals developed larger blisters and showed persistently reduced cellular infiltration compared with MHL. Interestingly, the MHO group displayed the most pronounced shift in blood flow between acute inflammation and resolution across all phenotypes. Collectively, these findings indicate distinct tissue and vascular inflammatory responses across metabolic groups.

### Blister exudate proteomics identifies phenotype-specific inflammatory dynamics

To define soluble inflammatory programs within the local blister response, we profiled the fluid fraction of blister exudates using a 96-plex Olink inflammation panel. We focused on proteins that differed across metabolic phenotypes, changed between acute inflammation and resolution, or contributed to group separation in PLS-DA analyses (**Fig. S2A**). Hierarchical clustering of these proteins revealed distinct temporal and phenotype-associated patterns (**Fig. 1E**).

Several proteins showed shared regulation across groups (Cluster 1), consistent with a core inflammatory response to cantharidin challenge. These included broadly engaged inflammatory, angiogenic, and tissue-stress mediators such as TNF, VEGFA, and IL-33 (Gundrathi et al., 2026; Pham and Kim, 2025; Zhu et al., 2024), indicating that major components of the blister response were preserved across metabolic phenotypes. In contrast, other protein clusters revealed marked phenotype specificity. Most notably, a resolution-associated cluster (Cluster 2) was induced from acute inflammation to resolution in MHL, MUL, and MHO individuals, but this induction was blunted in MUO. This cluster included IL-10, together with mediators linked to immune-cell recruitment, inflammatory polarization, and tissue remodeling, including CXCL1, CXCL10, IL-17A, IL-12B, and MMP10 (Bhaumik and Basu, 2017; Caley et al., 2015; Lee et al., 2013; Zhang et al., 2016). Thus, MUO individuals showed attenuated induction of a soluble mediator program associated with inflammatory resolution.

MUO individuals also showed a distinct acute-phase signature characterized by high levels of neutrophil-associated chemokines, including IL-8/CXCL8, CXCL5, and CXCL6 (Rajarathnam et al., 2019), together with tissue injury-associated mediators such as IL-1α and IL-22RA1 (Macleod et al., 2021; Wasserer et al., 2026) (Cluster 3). This pattern is consistent with an exaggerated early innate inflammatory response in MUO. This cluster also contained FGF-21, which was low in MHL but elevated particularly in MUO across both phases, suggesting that FGF-21 may reflect metabolic stress (BonDurant and Potthoff, 2018) rather than temporal inflammatory regulation alone. Additional clusters (Cluster 4, 5) showed reduced or attenuated mediator levels in metabolically unhealthy phenotypes, including immune activation and recruitment-associated proteins such as MCP-2, CD244, TNFSF14, TRAIL, and CCL4 (Chavez and Kiaris, 2025; Sindhu et al., 2019; Sun et al., 2021; Zheng et al., 2025), as well as tissue remodeling and type 2-associated mediators such as IL-13. Collectively, these data show that the local inflammatory proteome is dynamically shaped by metabolic phenotype, with MUO characterized by exaggerated early innate inflammation, elevated metabolic stress signals, and impaired induction or maintenance of mediators associated with resolution and tissue repair.

### Acute-phase blister mediators associate with clinical metabolic risk

To assess the clinical relevance of blister-derived inflammatory mediators, we focused on proteins with high Variable Importance in Projection (VIP) scores from PLS-DA analyses, as these features contributed most to separation between metabolic phenotypes. We selected the six proteins with the highest VIP scores from PLS-DA analyses done at three time points: the acute inflammation phase, the resolution phase, and the phase-to-phase change. (**Fig. S2A**). We then correlated these proteins with markers of systemic inflammation (CRP), glycemic control (fasting glucose, insulin, HOMA2-IR), and adiposity (sagittal height and waist circumference).

During the acute inflammation phase, several of these proteins correlated with clinical parameters of metabolic disease (**Fig. 1F, left panel**). FGF-21 showed significant correlations with glycemic control and adiposity, including HOMA2-IR (*r* = 0.57) and sagittal height (*r* = 0.49), but did not correlate with CRP. Consistent with this, blister FGF-21 levels were low in MHL but elevated across metabolically unhealthy and obese groups, but particularly in MUO (**Fig. 1E, cluster 3**). This pattern supports the interpretation that blister-derived FGF-21 reflects metabolic stress rather than temporal inflammatory regulation within the blister response or systemic inflammation as captured by CRP. Indeed, elevated circulating FGF-21 is a well-established feature of obesity and metabolic disease (Berti et al., 2015; Patt et al., 2024; Takebe et al., 2023), whereas its relationship with CRP is less consistent, with studies reporting either no association in obesity (Ebert et al., 2014), or positive associations in metabolic syndrome (Ebrahimi et al., 2021). Another notable finding during the acute inflammation phase was that several interleukins (IL-13, IL-33, IL-20) showed a negative correlation with CRP in the blister exudate, although IL-10 did not (**Fig. 1F**). This raises the possibility that early induction of type 2- and tissue repair–associated mediators may be more pronounced in individuals with lower systemic inflammation following cantharidin application, consistent with the role of these cytokines in barrier inflammation and repair responses described in atopic dermatitis (Hasegawa et al., 2022; Lu et al., 2022; Rojahn et al., 2020).

In contrast to the stronger associations observed during the acute inflammation phase, VIP-ranked proteins from the resolution phase showed fewer and less consistent correlations with clinical parameters **(Fig. 1F, middle panel)**. TNF and TRAIL correlated with measures of glycemic control, whereas uPA showed negative correlations with adiposity and systemic inflammation, including a moderate inverse correlation with CRP (*r* = -0.42). Similarly, phase-to-phase changes showed only limited associations with clinical parameters **(Fig. 1F, right panel)**, although adiposity measures such as sagittal height and waist circumference correlated negatively with changes in IL-8/CXCL8 (*r* = -0.46 and *r* = -0.42, respectively).

Collectively, these correlation analyses suggest that acute blister-derived inflammatory mediators more closely reflect underlying metabolic and adiposity-related alterations, whereas resolution-phase and dynamic change markers are less tightly linked to systemic clinical features.

### Transcriptomic profiling reveals impaired wound healing response in unhealthy metabolic phenotypes

We next performed RNA sequencing of blister-derived cells to assess transcriptional differences across metabolic phenotypes during acute inflammation (**Fig. 1G–J**). To evaluate how inflammatory responses changed over time, we compared phase-to-phase transcriptional changes between the acute and resolution phases within each phenotype. This enabled us to assess the dynamic regulation of the local inflammatory response and identify genes associated with the transition from acute inflammation to resolution across metabolic phenotypes.

Principal component analysis revealed that phase-dependent transcriptomic separation varied across metabolic phenotypes (**Fig. 1G**). MHL individuals displayed the clearest transcriptional shift between the acute and resolution phases, as reflected by separation in the PCA (**Fig. 1G**). Furthermore, the MHL group displayed a higher number of differentially expressed genes and more extensive pathway enrichment among significantly regulated genes with a greater effect size, defined as an absolute fold change greater than 0.5 (**Fig. 1H-I**). In comparison, the other phenotypes showed less pronounced transcriptional changes between phases, indicating that the transcriptional transition from acute inflammation to resolution was attenuated in these groups.

Across all metabolic phenotypes, the transition from inflammation to resolution was consistently associated with enrichment of transcriptional pathways related to immune cell chemotaxis (**Fig. 1H; Fig. S2B**) and included a shared reduction in acute neutrophil and inflammatory myeloid programs, including *CXCR1*, *CXCR2*, *FPR1*, *S100A12*, *TREM1*, and *IL1B* (Andrade et al., 2026; Dorward et al., 2015; Torfs et al., 2026), accompanied by increased expression of macrophage-associated clearance and remodeling genes such as *TREM2*, *GPNMB*, *FOLR2*, and *C1QA*/*C1QC* (Guan et al., 2025), together with tissue-repair factors *IGF1*, *HGF*, and *TIMP3* (**Fig. 1I**). These conserved changes define a common transcriptional backbone of the inflammatory-to-healing transition, characterized by attenuation of acute innate inflammation and engagement of clearance and tissue-restorative processes. These shared responses are consistent with the expected dynamics of the blister model, which captures coordinated changes in immune cell recruitment, tissue remodeling, and cellular composition during inflammatory resolution, as previously described by Gilroy and colleagues (Jenner et al., 2014).

Beyond these shared transcriptional responses, phenotype-specific programs distinguished the transition toward resolution (**Fig. 1I**). In MHL, this transition was characterized by coordinated suppression of innate inflammation and engagement of immune regulation and tissue restoration. Resolution was marked by suppression of neutrophil- and innate immune-associated genes, including *FCGR3B*, *CSF3R*, *TLR4*, *C5AR1*, and *IFNAR1* (Lyadova, 2025; Yan and Gao, 2012). This was accompanied by induction of adaptive and immunoregulatory genes, including *CXCL13*, *IL17F*, and *CD244* (Wang et al., 2023). Increased expression of *SIGLEC8* and *IL5RA* was paralleled by induction of matrix-remodeling genes *MMP11* and *MMP14* and epidermal differentiation genes *LCE1B* and *SPRR2A* (Gill and Parks, 2008; Hui et al., 2024; Meehan and Wang, 2022). Collectively, these changes supported engagement of reparative programs, consistent with progression toward immune regulation, remodeling, and barrier restoration (Yan et al., 2024).

In contrast, metabolically unhealthy phenotypes showed less coordinated engagement of inflammatory resolution and tissue repair. MUL showed induction of metabolic regulators including *PPARA*, *ACACB*, and *CYP27A1* (Lin et al., 2022), but reduced *IL33*, *IL13*, *F13A1*, and *IVL* (Tatu et al., 2022), suggesting metabolic adaptation without equivalent engagement of reparative and epithelial-restorative pathways. MUO displayed the most discordant signature, with increased *ELANE*, indicating persistence of neutrophil-associated activity (Wang et al., 2025), alongside increased *STAB1* and *LYVE1*, potentially reflecting a compensatory macrophage response (Guan et al., 2025). At the same time, *HBEGF*, *CASP14*, *TINCR*, and *PKP3* (Shirakata et al., 2005; Sun et al., 2015), as well as matrix-associated genes *FOXQ1* (Feuerborn et al., 2011) and *COL6A2* (Fitzgerald et al., 2013), were reduced, suggesting impaired epithelial differentiation and wound repair.

Notably, the relatively protected MHO phenotype was associated with a distinct transcriptional response combining inflammatory restraint with adaptation to hypoxic and metabolic stress. Reduced expression of inflammatory and lymphoid genes (*IL1A*, *ACOD1*, *CD8A*/*CD8B*, *FYN*) was accompanied by a tissue-preservation signature. This included non-inflammatory lipid handling (*HILPDA*) (van Dierendonck et al., 2022), chemokine clearance (*ACKR2*) (Gowhari Shabgah et al., 2022), hypoxic-mitochondrial quality control (*EGLN3*, *BNIP3*) (Irazoki et al., 2022), and microvascular matrix stabilization (*PDGFB*, *EPHB4*, *TIMP2*, *GUCY1A1*) (Liu et al., 2023).

Collectively, these findings suggest that metabolic health is associated with coordinated inflammatory resolution and tissue repair. This response was most evident in MHL, while MHO exhibited a distinct, potentially protective adaptive signature. In contrast, MUL and especially MUO showed less coordinated regulation of metabolic adaptation, inflammatory termination, and tissue restoration.

### Resolution-associated epithelial repair is attenuated in metabolically unhealthy phenotypes

To further assess phenotype-specific transcriptional dynamics, we performed enrichment analyses and compared the phase-to-phase (72 vs 24h) log2 fold change for each gene between metabolically healthy and unhealthy phenotypes within the lean and obese groups **(Fig. 1J)**. These plots allow direct comparison of phase-to-phase regulation between phenotypes, with concordant or divergent gene positioning indicating shared or phenotype-specific transcriptional responses, respectively. This analysis revealed that several epidermal differentiation and barrier-associated genes, including late cornified envelope (CE) family members, were induced during resolution in the metabolically healthy groups (MHL and MHO) **(Fig. 1J)**, consistent with activation of tissue repair programs. The CE forms the outermost layer of the skin and exhibits barrier, antioxidant and antimicrobial functions (Candi et al., 2005; Henry et al., 2012). Upregulation of structural CE genes, such as small proline-rich proteins (SPRRs, like *SPRR2F* and *SPRR3*), or late cornified envelope genes (*LCE1A-B*, *LCE2A-D*) in metabolically healthy individuals indicates proper re-establishment of the CE in the skin blisters, indicative of tissue repair and wound healing as part of the resolution program. However, participants in the MUL and MUO groups fail or show a delay in engaging the gene program required for CE formation, suggesting a link between metabolic health and cutaneous disorders. Indeed, metabolic syndrome has been linked to increased instances of several skin disorders, such as psoriasis, acne and hidradenitis suppurativa (Elzawawi et al., 2024; Mitamura et al., 2021; Steele et al., 2019), which may be reflected in our blister model. Indeed, in addition to the phenotype-specific differences in CE remodeling, a common denominator across groups was that the phase-to-phase transition was consistently associated with enrichment of pathways related to epidermal and skin development, as well as keratinocyte biology. This is expected, as the blister model captures not only leukocyte infiltration but also dynamic changes in the local skin environment. Consistent with this, previous studies have shown that blister exudate samples contain keratinocytes, melanocytes, fibroblasts, and some smooth muscle cells, in addition to leukocytes (Rojahn et al., 2020).

### Flow cytometry analysis of blister-derived cells reveals distinct immune cell composition and activation across metabolic phenotypes

To complement the transcriptomic analysis with direct assessment of immune cell composition and surface marker expression, we also performed flow cytometry on blister-derived leukocytes collected during acute inflammation and resolution phase, respectively (**Table 2**; gating strategy in **Fig. S3A–B**).

**Table 2:**
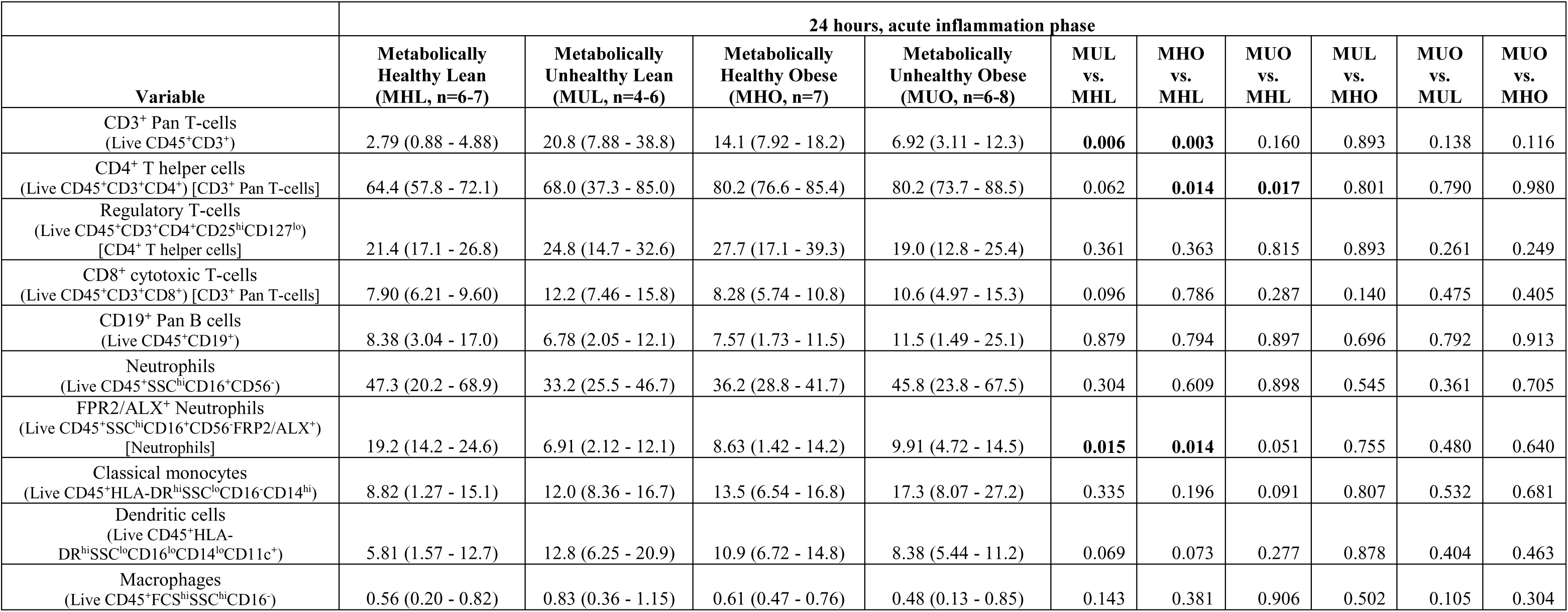
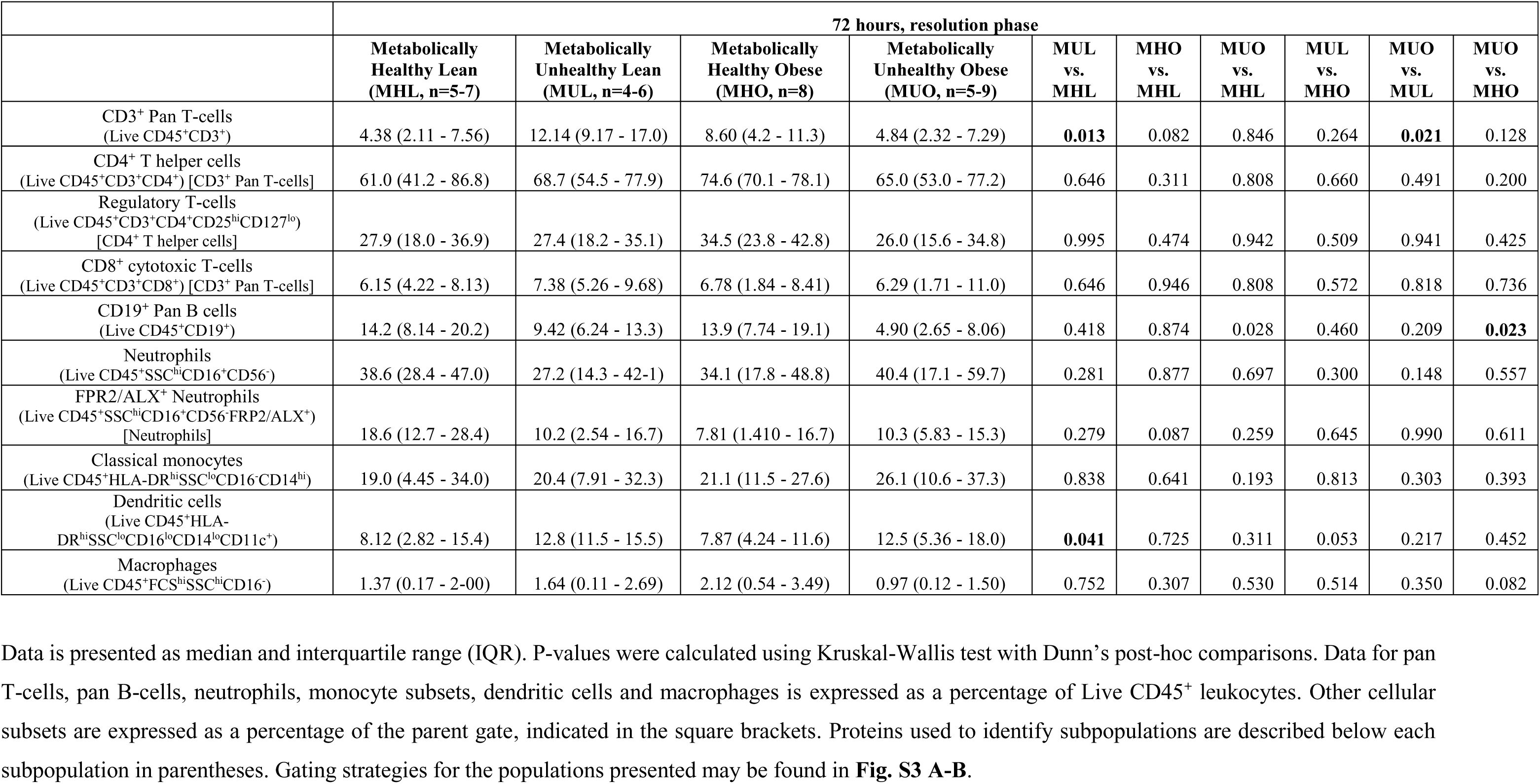
Immunophenotyping of blister exudate leukocytes. Data is presented as median and interquartile range (IQR). P-values were calculated using Kruskal-Wallis test with Dunn’s post-hoc comparisons. Data for pan T-cells, pan B-cells, neutrophils, monocyte subsets, dendritic cells and macrophages is expressed as a percentage of Live CD45^+^ leukocytes. Other cellular subsets are expressed as a percentage of the parent gate, indicated in the square brackets. Proteins used to identify subpopulations are described below each subpopulation in parentheses. Gating strategies for the populations presented may be found in **Fig. S3 A-B**.

A distinct immunological feature of the MUL group was a higher frequency of CD3^+^ pan-T cells. During the acute phase, CD3^+^ pan-T cell frequencies were increased in MUL individuals compared with MHL controls (MUL, 20.8% [7.88–38.8] vs. MHL, 2.79% [0.88–4.88]; p = 0.006). A similar, although less pronounced, increase was observed in the MHO group during acute inflammation, but this difference was not sustained during the resolution phase. These findings suggest that T cells make up a larger proportion of the recruited inflammatory-cell compartment in MUL individuals compared with MHL controls. The persistence of this relative T cell enrichment into the resolution phase may reflect prolonged T cell retention, delayed clearance, or sustained immune surveillance at the site of tissue injury. However, because CD3 identifies total T cells, these data do not define which T cell subsets are involved. Nevertheless, this T cell-enriched profile supports the concept that MUL may represent a lean but immunometabolically activated phenotype, in which metabolic disease is associated with local immune dysregulation independent of adiposity.

We next assessed myeloid-cell subsets in blister fluid across metabolic phenotypes. Classical CD16^-^CD14^hi^ monocytes cells were detected in the exudates but did not differ between phenotypes during either the acute or resolution phase. Intermediate and non-classical monocytes were detected at very low frequencies and were therefore excluded from further analysis. Macrophage frequencies, defined as live FSC^hi^SSC^hi^CD16^+^ cells, also did not differ between phenotypes at either time point. In contrast, dendritic cells, defined as live CD16^lo^CD14^lo^CD11c^+^ cells, were increased in MUL individuals compared with MHL controls during both acute inflammation and resolution. This difference approached statistical significance during the acute phase (MUL, 12.8% [6.25–20.9] vs. MHL, 5.81% [1.57–12.7]; p = 0.069) and reached significance during resolution (MUL, 12.8% [11.5–15.5] vs. MHL, 8.12% [2.82–15.4]; p = 0.041). Together with the increased frequency of CD3^+^ pan-T cells in MUL blister fluid, these data suggest that the MUL phenotype is characterized by a coordinated enrichment of antigen-presenting and T cell compartments within the local inflammatory infiltrate, supporting the concept of immune dysregulation in metabolically unhealthy lean individuals.

B cell frequencies did not differ markedly between metabolic phenotypes during the acute inflammation phase. However, during resolution, MHL individuals showed a robust increase in CD19^+^ B cells among blister-fluid cells, whereas this response was absent in MUO individuals. Consequently, MUO individuals had a lower frequency of CD19^+^ B cells during the resolution phase compared with both MHL and MHO individuals. This suggests that MUO is associated with impaired resolution-phase accumulation of B cells at the site of tissue injury, potentially reflecting altered recruitment or retention of cells involved in antigen presentation, immune regulation, and tissue homeostasis (Oleinika et al., 2022).

We next assessed neutrophil numbers, which did not differ between phenotypes. To determine whether metabolic phenotype was associated with altered resolution signaling, we examined neutrophil expression of FPR2/ALX, the receptor for the specialized pro-resolving mediator lipoxin A4, which is implicated in the regulation of inflammatory resolution. (Chiang and Serhan, 2020; Serhan et al., 2007). During acute inflammation, FPR2/ALX expression was lower in MUL, MHO, and MUO individuals than in MHL controls, reaching statistical significance in MUL and MHO individuals and approaching significance in MUO individuals (MUL, p = 0.015; MHO, p = 0.015; MUO, p = 0.051; **Table 2**). A similar pattern was observed during the resolution phase, although these differences did not reach statistical significance **(Table 2**). These findings suggest that altered adiposity and metabolic dysfunction are associated with reduced acute expression of a key pro-resolving receptor in the local inflammatory response.

### Systemic inflammatory markers only modestly distinguish metabolic phenotypes

To evaluate systemic inflammation across metabolic phenotypes, we combined routine clinical inflammatory markers, circulating leukocyte phenotyping, and transcriptomic profiling of blood-derived cells (**Fig. 2A**). We first assessed conventional inflammatory parameters from the clinical biochemistry panel (**Table 1**). Consistent with previous studies (Esser et al., 2014; Su et al., 2024), circulating markers of low-grade inflammation, including CRP, were elevated in individuals with obesity compared with lean individuals. However, these markers showed limited ability to distinguish metabolic health status within obesity, with no clear differences between MHO and MUO individuals. We next examined routinely measured circulating leukocyte populations. Neutrophil counts were elevated in MUO individuals compared with MHL controls, whereas this pattern was not observed in the other phenotype comparisons. Eosinophil counts differed between MHO and MHL individuals, but absolute values were low, and this finding was therefore interpreted cautiously. Across clinical inflammatory measures, no major differences were observed between MHL and MUL individuals, indicating that routine systemic inflammatory markers did not clearly capture the metabolically unhealthy lean phenotype.

**Figure 2.**
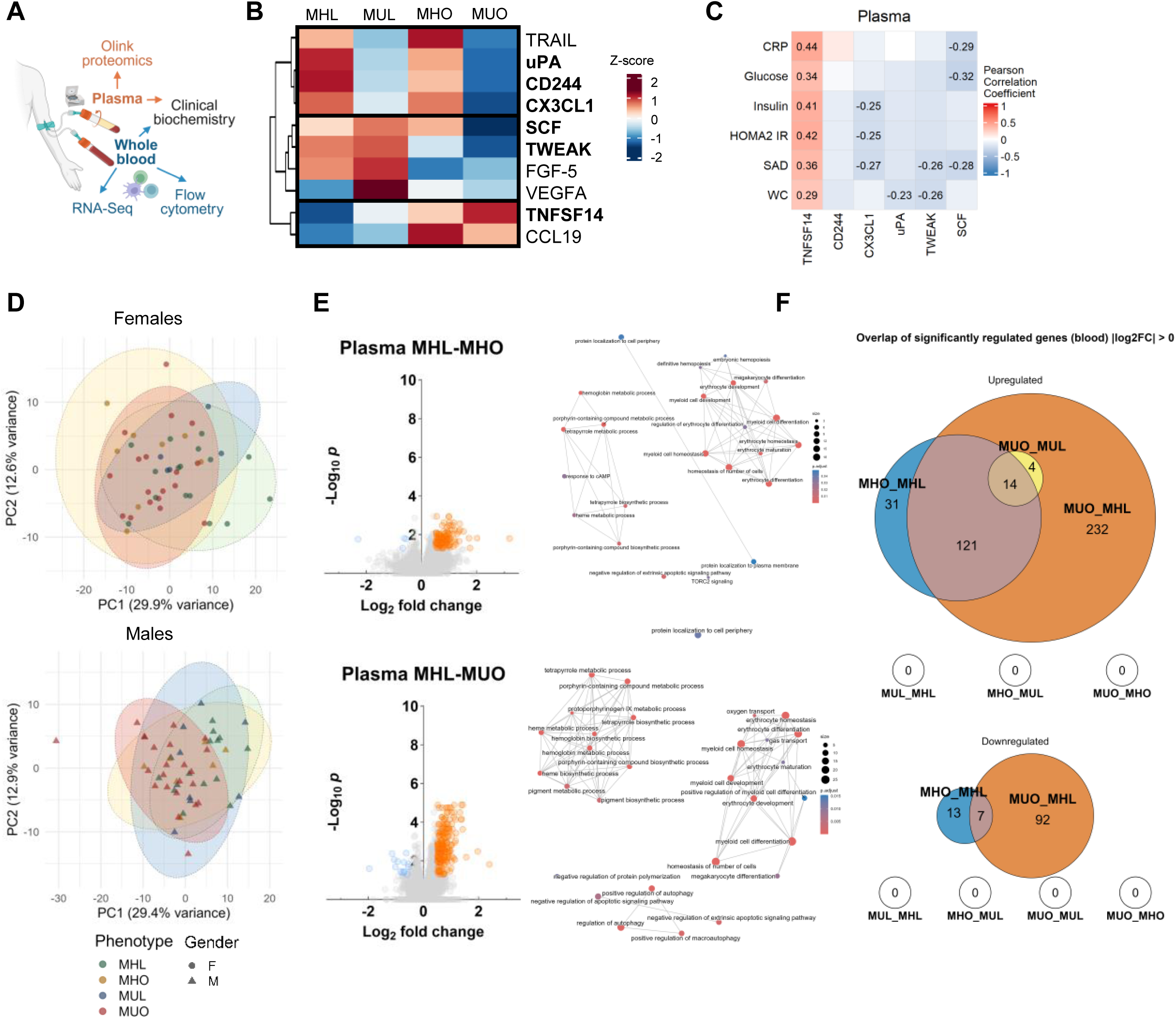
Plasma protein expression levels and RNA-Seq analysis related to inflammatory processes. **(A)** Graphical representation of the blood samples collected for systemic immunophenotyping, plasma Olink proteomics, and RNA sequencing of blood derived cells. **(B)** Heatmap and hierarchical clustering of significantly different plasma protein expression levels related to inflammatory processes measured using the Olink Inflammation panel. Data are presented as row-wise scaled and centered means (Z-score) of NPX values in log2 scale. One-way ANOVA with Tukey’s HSD post-hoc test was used and p ≤ 0.05 was considered statistically significant. **(C)** Heatmap with Pearson correlation coefficients (*r*) between the top-ranking plasma proteins, identified by VIP scores from the PLS-DA model, and log2-transformed clinical markers of systemic inflammation (CRP), glycemic control (fasting glucose, insulin, HOMA2-IR), and adiposity (sagittal height, waist circumference). The color scale represents the strength and direction of the correlation (red = positive, blue = negative). Correlation coefficients are numerically displayed only for statistically significant associations (p ≤ 0.05). **(D)** Principal component (PCA) analysis of blood cell-derived transcriptome across metabolic phenotypes based on the top 500 genes with the highest inter-sample variance. Axis labels indicate the percentage of total variance explained by each component. **(E)** Volcano plots of RNA sequencing data comparing MHL to MHO (top panel) and MHL to MUO (bottom panel). **(F)** Euler diagrams showing the overlap of significantly upregulated (top) and downregulated (bottom) genes in blood derived cells across phenotype comparisons. MHL, Metabolically Healthy Lean; MUL, Metabolically Unhealthy Lean; MHO, Metabolically Healthy Obese; MUO, Metabolically Unhealthy Obese.

### Circulating immune-cell profiling reveals selective T-cell differences across metabolic phenotypes

To complement routine clinical immunophenotyping, we performed more detailed flow cytometric profiling of circulating leukocytes. Differences across metabolic phenotypes were largely confined to the T-cell compartment and were most evident in MHO (**Table 3, Fig. S3C–E**).

**Table 3:**
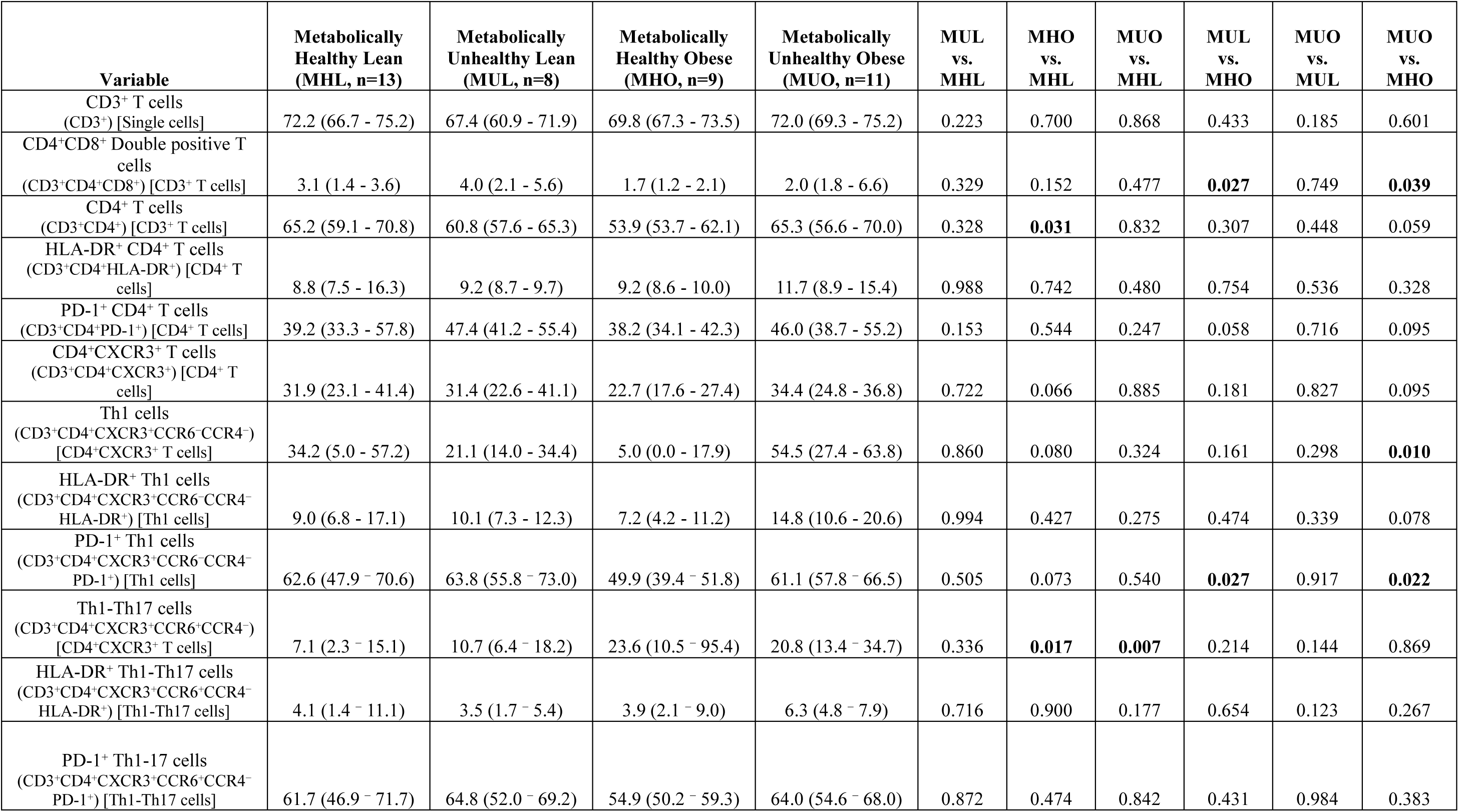
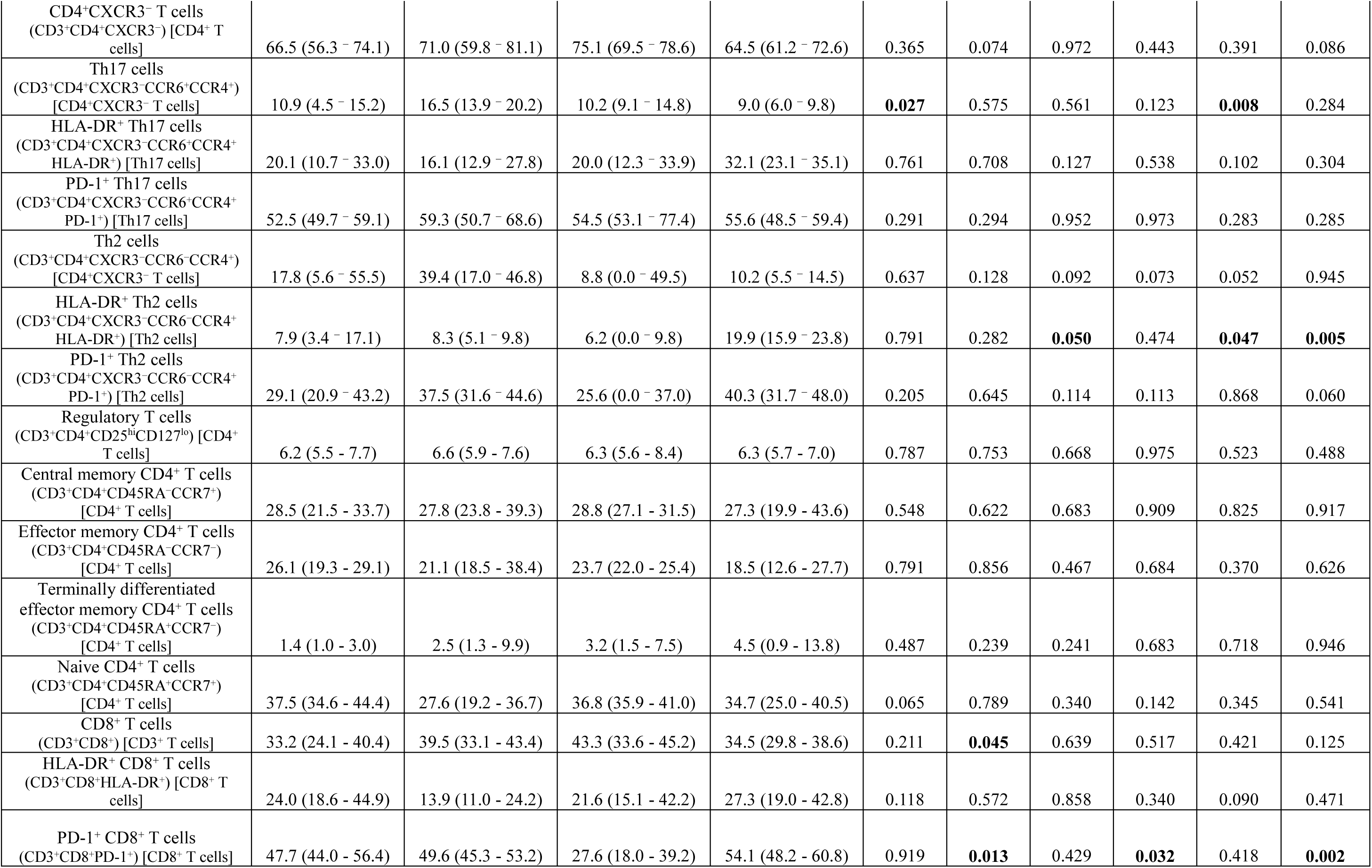
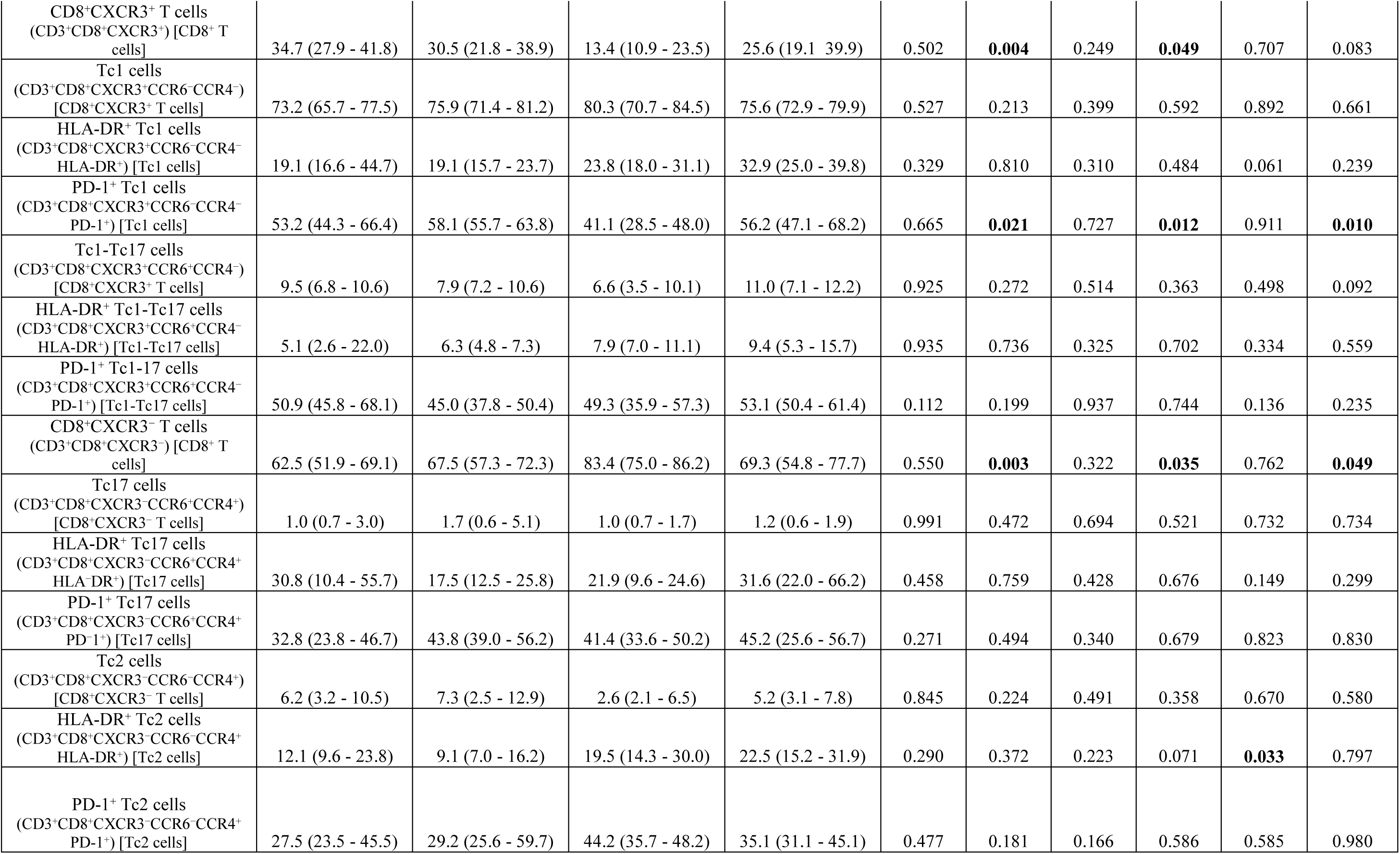
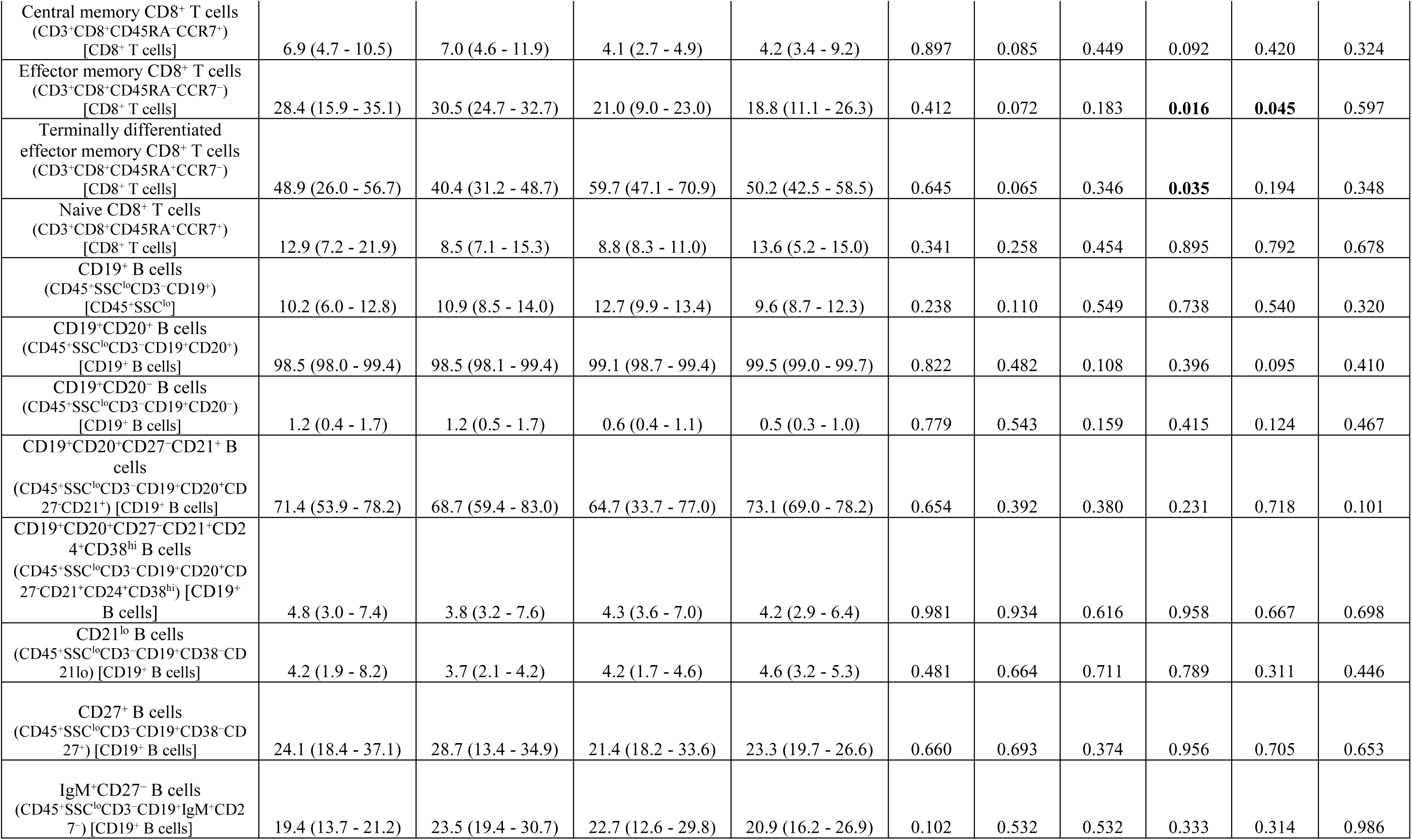
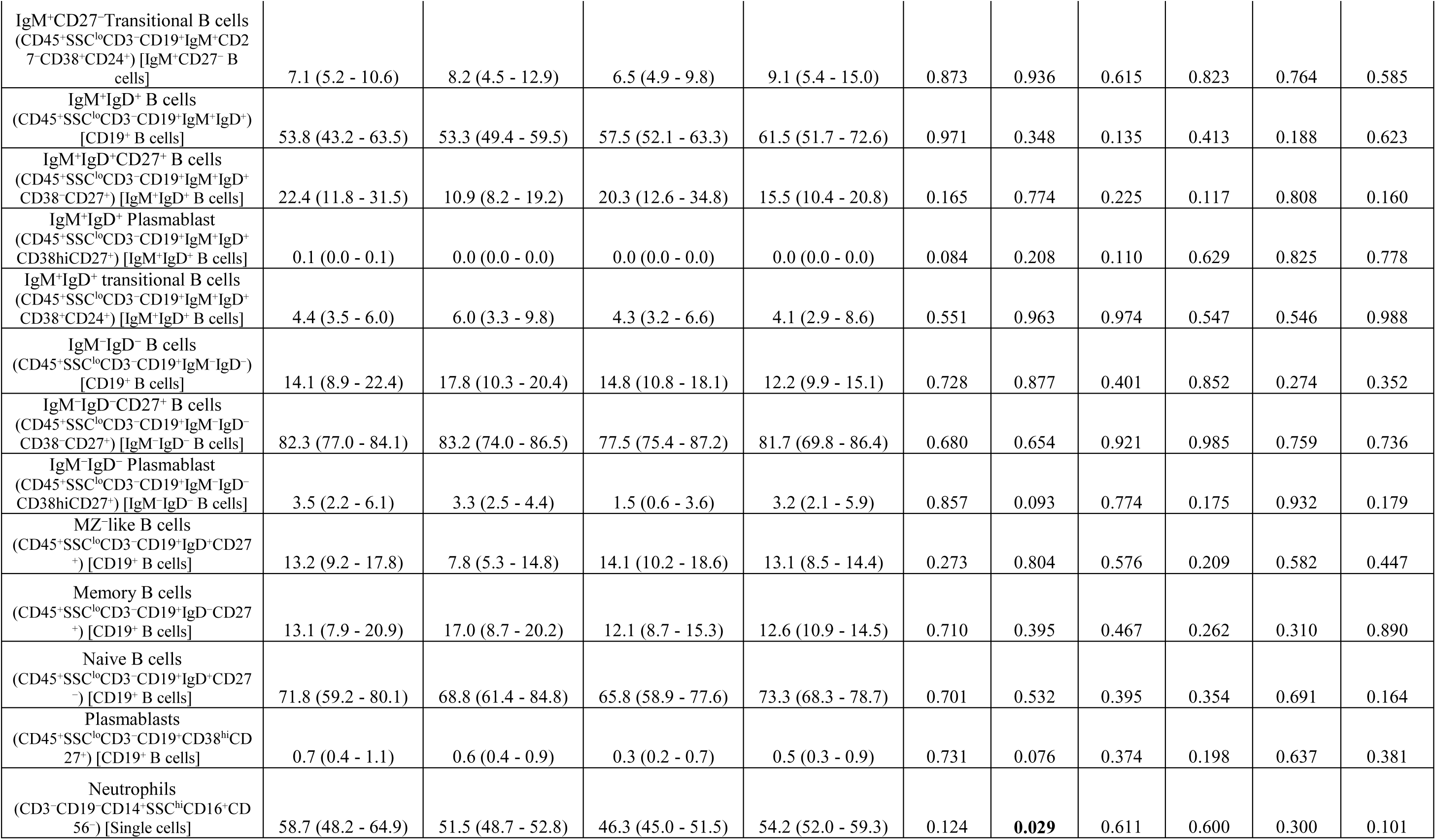
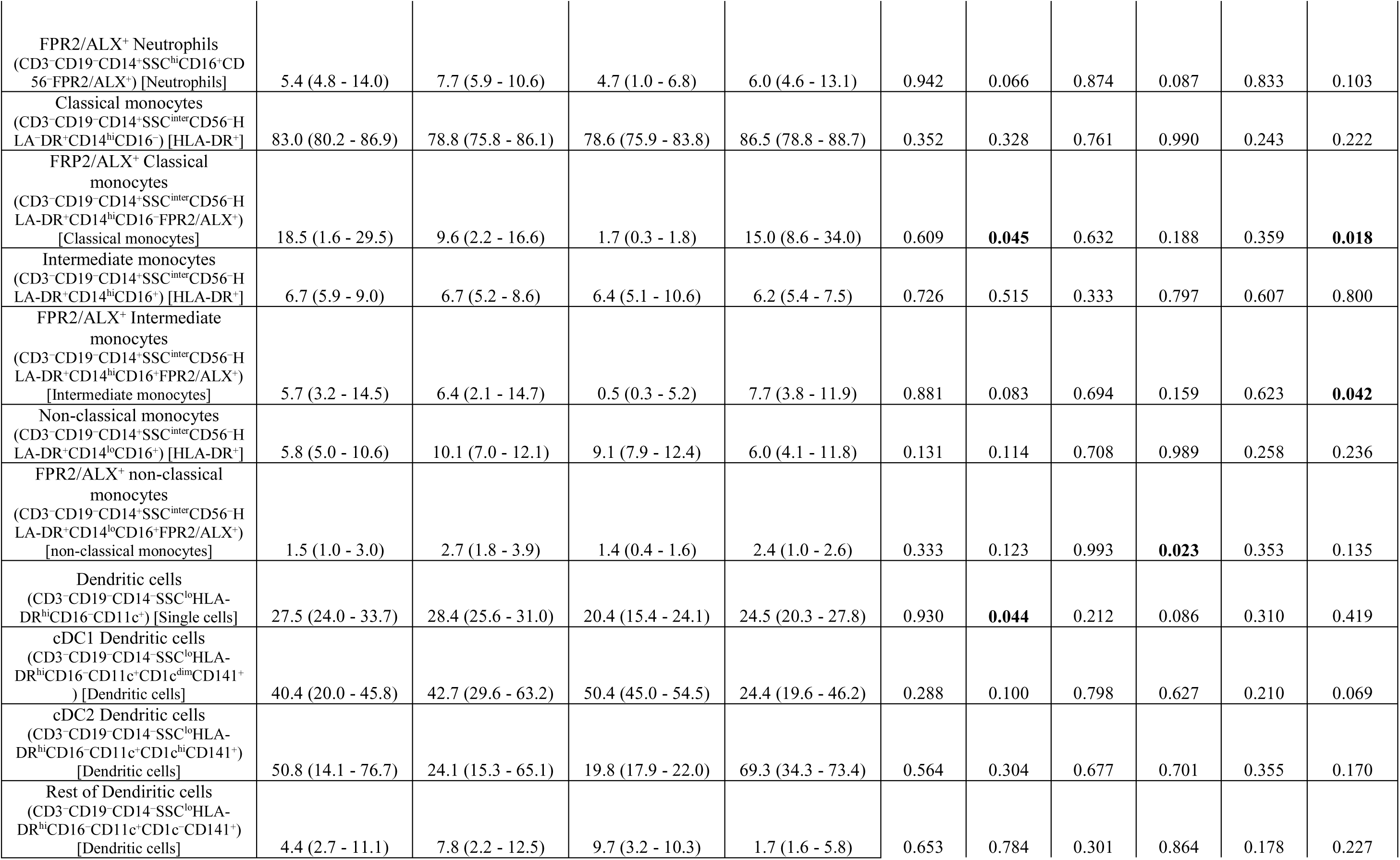
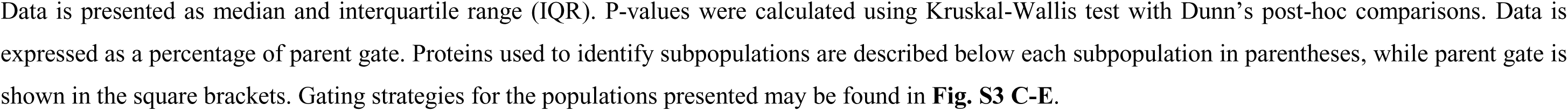
Immunophenotypes of circulating leukocytes. Data is presented as median and interquartile range (IQR). P-values were calculated using Kruskal-Wallis test with Dunn’s post-hoc comparisons. Data is expressed as a percentage of parent gate. Proteins used to identify subpopulations are described below each subpopulation in parentheses, while parent gate is shown in the square brackets. Gating strategies for the populations presented may be found in **Fig. S3 C-E**.

MHO individuals had lower frequencies of circulating CD4^+^ T cells than MHL individuals, accompanied by increased CD8^+^ T-cell frequencies (**Table 3**). CD4^+^CD8^+^ double-positive T cells were also reduced in MHO compared with metabolically unhealthy individuals (**Table 3**). PD-1 expression on CD8^+^ T cells was lower in MHO than in all other phenotypes (**Table 3**). Given the association of PD-1 with chronic immune activation and T-cell exhaustion (Jubel et al., 2020), this pattern is consistent with altered T-cell activation in MHO, although PD-1 expression alone does not establish exhaustion.

By contrast, circulating B-cell subsets and monocytes did not differ between metabolically healthy and unhealthy groups (**Tables 1 and 3**). Neutrophil levels were modestly increased with obesity. We also assessed FPR2/ALX expression as a marker of resolution-related receptor availability. Neutrophil FPR2/ALX expression was numerically lower in MHO than in MUO, but this difference was not statistically significant (**Table 3**).

Collectively, baseline circulating immune profiles showed limited differences across metabolic phenotypes, with selective changes in T-cell composition and PD-1 expression in MHO. The greater heterogeneity observed in the skin-blister model suggests that dynamic tissue responses reveal differences in immune regulation that are less apparent in steady-state blood measurements.

### Circulating inflammatory biomarkers show modest phenotype-associated variation

To determine whether inflammatory differences observed in the blister exudate were reflected systemically, we performed plasma proteomic profiling using the Olink inflammation panel **(Fig. 2B–F)**. In contrast to the pronounced phenotype- and phase-associated patterns observed in blister fluid, plasma inflammatory profiles showed only limited separation between metabolic phenotypes. Only 10 of the 92 assessed inflammatory markers differed significantly across groups, indicating relatively modest systemic inflammatory divergence **(Fig. 2B)**.

Among the differentially regulated proteins, some markers were consistent with metabolic disease and adiposity-associated inflammation. Cluster 1 included uPA, which has recently been implicated in metabolic dysfunction in a mouse model of diet-induced obesity, where uPA-deficient mice were initially protected from weight gain and metabolic impairment, although prolonged high-fat diet exposure eventually led to metabolic dysfunction (Hur et al., 2026). Clusters 2 and 3 contained proteins associated with adiposity, including SCF and TWEAK, which were reduced in individuals with obesity. Although circulating SCF levels were reduced in the present study, previous work has linked reduced cutaneous SCF to delayed wound closure in models of aging, obesity, and alcohol exposure (Wang et al., 2020). Conversely, TNFSF14 was increased, particularly in MUO individuals compared with MHL controls. Together, these patterns suggest that most systemic biomarker variation was driven by adiposity rather than metabolic health status per se. This interpretation is consistent with previous studies linking elevated TNFSF14 to Prader-Willi syndrome (Faienza et al., 2023), childhood obesity (Faienza et al., 2019), and other obesity-associated conditions (Bassols et al., 2010; Halvorsen et al., 2016).

### Circulating inflammatory biomarkers show limited association with clinical metabolic traits

To assess the clinical relevance of the plasma proteins contributing most strongly to phenotype separation, we selected the six proteins with the highest VIP scores from the PLS-DA analysis **(Fig. S4)** and correlated their circulating levels with markers of systemic inflammation, glycemic control, and adiposity **(Fig. 2C)**. Despite being the top contributor to group separation, uPA showed only weak associations with clinical variables, including a modest inverse correlation with waist circumference (*r* = −0.23), and no significant correlations with the remaining parameters. Similarly, CD244, CX3CL1, and TWEAK did not correlate significantly with CRP, indicating that these markers are not captured by conventional measures of systemic inflammation. SCF showed a statistically significant but modest inverse correlation with CRP (*r* = −0.29). In contrast, TNFSF14 displayed the strongest and most consistent associations, including a positive correlation with CRP (*r* = 0.44) and broader associations with metabolic and anthropometric measures. Together, these findings identify TNFSF14 as a marker of adiposity-associated systemic inflammation rather than metabolic health status, as TNFSF14 levels were comparable between MHL and MUL individuals and between MHO and MUO individuals.

### Whole-blood transcriptomics provides limited separation of metabolic phenotypes

To determine whether circulating immune-cell transcriptional profiles provided greater resolution of systemic inflammatory differences, we next performed whole-blood RNA sequencing **(Fig. 2D–F)**. Principal component analysis revealed limited separation between metabolic phenotypes, indicating that global transcriptional profiles in circulating blood cells did not clearly distinguish the four groups **(Fig. 2D)**. Differential expression and pathway analyses showed modest differences between lean and obese individuals, but provided limited discrimination between metabolically healthy and metabolically unhealthy phenotypes within either adiposity group **(Fig. 2E–F)**. Thus, consistent with the plasma biomarker analyses, whole-blood transcriptomics captured broad adiposity-associated differences but did not identify a distinct systemic transcriptional signature of metabolic health status. Collectively, these findings indicate that circulating inflammatory biomarkers and blood-cell transcriptional profiling have limited utility for resolving metabolic phenotypes compared with the more pronounced phenotype-specific inflammatory programs observed in blister exudates.

## Concluding remarks

In this study, we integrated systemic immune profiling with a human *in vivo* model of peripheral tissue inflammation to examine inflammatory regulation across metabolic phenotypes. While low-grade systemic inflammation was elevated in obesity, consistent with previous reports, differences between metabolically healthy and unhealthy individuals were only modest. Plasma proteomic profiling identified a limited set of discriminatory proteins, and profiling of circulating leukocytes as well as transcriptional analysis of whole blood revealed minimal differences between phenotypes, indicating that steady-state systemic measurements provide limited resolution to distinguish metabolic health status.

In contrast, peripheral tissue inflammation revealed pronounced and coordinated differences across proteomic, cellular, and transcriptomic layers. Proteomic profiling of blister exudates demonstrated distinct temporal inflammatory trajectories, with metabolically unhealthy individuals exhibiting exaggerated early innate responses together with impaired engagement of resolution- and tissue repair–associated pathways. These findings were supported by flow cytometry analysis, which showed altered immune cell composition and turnover of blister leukocytes, including reduced accumulation of reparative cell populations and sustained T-cell presence in metabolically unhealthy states.

At the transcriptional level, dynamic analysis of blister-derived cells between inflammation and resolving phases further highlighted divergent gene programs. Although engagement of inflammatory signaling and skin development was observed across all phenotypes, metabolically healthy individuals showed more significant and extensive activation of these gene programs compared to metabolically unhealthy groups. Metabolically unhealthy groups displayed specifically strong impairment of cornified envelope formation genes in the blister model, which may be linked to increased instances of skin disorders in metabolic syndrome patients.

The findings in this study should be interpreted in the context of cohort recruitment. Re-recruitment from the Scandinavian cohorts yielded participants with relatively early metabolic perturbations and limited medication use, rather than substantial insulin resistance or advanced cardiometabolic disease. This allowed us to identify local immune dysregulation in the setting of relatively modest systemic alterations, but may limit generalizability to more advanced disease. Studies spanning a broader range of disease severity, including longitudinal follow-up, will be needed to determine how these tissue-level immune alterations relate to cardiometabolic progression.

Collectively, these findings demonstrate that metabolic health is more accurately reflected in the regulation and kinetics of peripheral tissue inflammation than in baseline systemic immune parameters. The dissociation between systemic and tissue-level responses suggests that impaired transition from acute inflammation to resolution may represent a key feature of metabolic dysfunction. These results highlight the importance of dynamic *in vivo* models to uncover immunological heterogeneity and provide a framework for understanding how altered inflammatory regulation contributes to cardiometabolic disease.

## Acknowledgements

The work of E.B. is supported by the European Research Council (ERC-StG no. 804418), Independent Research Fund Denmark (DFF #3165-00026B), Carlsberg Foundation (CF24-2226) Aarhus University Research Foundation (AUFF-E-2022-7-8), Novo Nordisk Foundation (NNF22OC0079363), the Swedish state’s ALF-agreement (ALFGBG-978978), Regionala FoU-medel, Västra Götalandsregionen (OLG-2023-02-22) and the Swedish Research council (VR 2023-02627). Support for the work of S.L. comes from NIH (HL152251, HL128457), the Novo Nordisk Foundation (NNF22OC0079368), the Aarhus University Research Foundation (AUFF-E-2022-7-9), the Lundbeck Foundation (R396-2022-189), Independent Research Fund Denmark (DFF, #3165-00028B) and the CAPTURE HFpEF initiative. Technical assistance is acknowledged from Negar Ayoubzadeh. Flow cytometry was performed by staff at the flow cytometry unit at the Department of Clinical Immunology, Linköping University Hospital, Sweden, and at the FACS Core Facility, Aarhus University, Denmark. SCAPIS is supported by the Swedish Heart and Lung Foundation, the Knut and Alice Wallenberg Foundation, the Swedish Research Council and VINNOVA. We express our sincere gratitude to all study participants, with special thanks extended to the test personnel at the SCAPIS test center in Gothenburg.

## Author contributions

S. Tavajoh: data curation, investigation, methodology, validation, and writing – original draft, review and editing. B.E. Suur: data curation, formal analysis, investigation, validation, visualization, and writing – original draft, review and editing. M. Clark: investigation, writing – review and editing. A.A. Becerril-Campos: data curation, investigation, validation, visualization, and writing – original draft, review and editing. E. Velasco: investigation, writing – review and editing. B. Medelyte: investigation, writing – review and editing. L.B. Nørregaard: investigation, writing – review and editing. S.J. Tingskov: investigation, writing – review and editing. A. Bæk: investigation, writing – review and editing. L. Skaarup: investigation, writing – review and editing. E. Schrøder: investigation, writing – review and editing. M. Dost: investigation, writing – review and editing. S.B. Gribsholt: writing – review and editing. J.M. Bruun: writing – review and editing. M.Q. Järbrink: writing – review and editing. S. Nyström: methodology, writing – review and editing. I. Bergström: investigation, methodology, writing – review and editing. C. Åhlund: data curation, investigation, project administration, writing – review and editing. J.P. Pedersen: writing – review and editing. N. Jessen: writing – review and editing. L. Lin: methodology, resources, writing – review and editing. A. Harazin: investigation, validation, project administration, resources, writing – review and editing. H.H. Thomsen: writing – review and editing. P.A. Jansson: writing – review and editing. R. Blomgran: methodology, conceptualization, data curation, writing – review and editing. S. Lange: conceptualization, formal analysis, validation, and writing – original draft, review and editing. M. Soták: conceptualization, data curation, formal analysis, investigation, methodology, project administration, resources, software, supervision, validation, visualization, and writing – review and editing. E. Börgeson: conceptualization, funding acquisition, investigation, project administration, resources, supervision, validation, and writing – original draft, review and editing.

## Declaration of generative AI and AI-assisted technologies

ChatGPT was utilized for language editing to improve readability and clarity of the text in this publication.

## SUPPLEMENTAL FIGURE LEDGENDS

### Materials and Methods

#### Study design and human subjects

Participants were recruited from the Swedish CArdioPulmonary BioImage Study (SCAPIS) and the Danish Health in Central Denmark (HICD) cohorts (Bergström et al., 2015; Riis et al., 2020), as outlined in the flowchart (**Fig. S1)**. SCAPIS candidates were pre-stratified in 2020 by BMI and metabolic health phenotype. To establish a balanced design based on the smallest subgroup (metabolically unhealthy men, n = 57), age-matched SCAPIS participants were selected across sexes and phenotypic groups (n = 60 per group), prioritizing medication-free individuals prior to formal invitation. Concurrently, HICD participants were randomly selected for invitation from the pool of individuals who had provided prior consent for future contact. Adults eligible for inclusion underwent screening and were stratified according to predefined metabolic and adiposity criteria, as outlined below. Exclusion criteria were smoking, use of anti-inflammatory or immunosuppressive medication, chronic inflammatory disease, ongoing infection, cancer under treatment, significant gastrointestinal or inflammatory bowel disease, bleeding disorders, anticoagulant use, and recent alcohol intake.

All participants provided written informed consent, and the study was conducted in accordance with the Declaration of Helsinki. The study was approved by the Swedish Ethical Review Authority (ID: 2019-04179) and the Danish Scientific Ethics Committee (ID: 1-10-72-102-23) and was registered at ClinicalTrials.gov (NCT04256330 and NCT06390189).

#### Clinical assessment and participant categorization

Participants underwent a comprehensive health examination and medical history assessment. This included anthropometric measurements (e.g. BMI, waist, hip, neck and arm circumferences), vital signs (e.g. body temperature, blood pressure and heart rate), capillary blood glucose level, and blood collection for lipid profile and glycated hemoglobin (HbA1c) analysis. For HbA1c analysis, capillary blood was collected in a cartridge (DCA HbA1c Reagent Kit, Siemens, Cat#10698915) and analyzed in a semi-automated bench analyzer (DCA Vantage, Siemens).

In this cross-sectional observational study, all participants were stratified into four distinctive groups based on their body mass index (BMI) and metabolic health. BMI categories were lean (BMI < 25 kg/m²) and obese (BMI ≥ 30 kg/m²). Following criteria were used for patient stratification: I) elevated waist circumference (women ≥80 cm, men ≥ 94 cm), II) increased HbA1c (≥ 39 mmol/mol), or glucose-lowering medication use, III) increased triglycerides (≥ 1.7 mmol/L), or lipid-lowering medication use, IV) low HDL-Cholesterol (women < 1.29 mmol/L, men < 1.03 mmol/L) or anti-hyperlipidemia medication use, V) hypertension (systolic > 130 mmHg or diastolic > 85 mmHg), or anti-hypertensive medication. Participants presenting with 0–2 criteria were categorized as metabolically healthy, whereas those meeting ≥ 3 criteria were stratified as metabolically unhealthy, resulting in MHL, MUL, MHO and MUO groups.

#### Acute tissue inflammation-resolution assessment

A cantharidin-induced skin blister model was utilized on a subset of patients (n = 48; 5-6 per metabolic group and per sex) to characterize the recruitment and dynamics of infiltrating immune cells during the acute and resolving phases of inflammation (Jenner and Gilroy, 2012). To obtain sufficient material for subsequent analyses, four blisters were induced on each forearm at staggered intervals to capture distinct inflammatory phases. 12.5 µL of cantharidin 0.1% (Dormer Laboratories, Cat#9001-975M) diluted in acetone (Merck, Cat#1000220500), was applied to 10 mm filter paper discs using a Hamilton syringe (Merck, Cat#20735). The cantharidin-containing disc was secured under occlusive bandaging until harvest of the acute inflammation phase (dominant arm) 24 h post cantharidin challenge. The resolution phase (non-dominant arm) utilized a 24 h cantharidin-induction followed by a 48h recovery period with the blister rebandaged under a ventilated protective dressing, totaling 72h from the initial insult to sample collection. We quantified blister fluid volume across the time points, by weighing the collected tubes on a microgravity scale. As expected, based on the design of the model, fluid volume was low during the acute phase, when the cantharidin-containing disc is secured under occlusive bandaging, and increased during the resolving phase, when the blister is allowed to progress under a ventilated protective dressing. However, fluid volume did not differ across metabolic phenotypes, indicating comparable conditions between phenotypes.

#### Clinical chemistry

Venous blood was collected after an overnight fast and analyzed in the hospital’s accredited clinical chemistry laboratory. Citrate tubes (BD, Cat#362782) were used for coagulation assays (APTT, PK). Fluoride–citrate (FC) mix tubes (Greiner Bio-One, Vacuette, Cat#454513) were used for glucose. K₂EDTA tubes (Greiner Bio-One, Vacuette, Cat#454410) were used for erythrocyte sedimentation rate, platelet count, and leukocyte count. Lithium heparin tubes (BD, Cat#368497) were used for liver function (ALT, AST, ALP, albumin, protein, GGT, bilirubin), kidney function (creatinine, eGFR, sodium, potassium, calcium), and inflammatory status (CRP). All analytes are summarized in **Table 1**.

#### Saliva analyses

Morning and evening saliva samples were collected for cortisol measurement. Patients self-collected samples using Salivette tubes (Sarstedt, Cat#51.1534) according to the manufacturer’s instructions, avoiding exercise for 1.5 h and fasting for 1 h before collection. Samples with visible discoloration or sediment and those from individuals receiving cortisol treatment, with circadian rhythm disturbances, or tobacco use (oral nicotine pouches) were excluded. Tubes were centrifuged for 2 min at room temperature (RT) at 1000 × g (EBA 200, Hettich). Cortisol was quantified by ELISA (R&D Systems, Cat#KGE008B) after 1:4 dilution of saliva samples.

#### Urine analyses

Participants collected first-morning urine in sterile cups for the evaluation of early markers of renal dysfunction associated with cardiometabolic risk. Samples were centrifuged at 3000 rpm for 10 min (EBA 200, Hettich) at RT, aliquoted, snap frozen in liquid nitrogen and stored at −80°C until analysis. Urinary microalbumin and creatinine were quantified using the DCA Microalbumin/Creatinine (ACR) urine test (Siemens, Cat#01443699 6011A) on a semi-automated bench analyzer (DCA Vantage, Siemens) according to the manufacturer’s instructions.

#### Blister exudate collection and processing

At 24h (“Inflammation” blister) and 72h (“Resolution” blister) post-induction, bandages were removed and blister diameters measured. 20 min after bandage removal, blisters were punctured with a sterile needle. Fluid was carefully aspirated with sterile tip and transferred into 50 µL of 3% citrate buffer in saline (Sigma-Aldrich, Cat#1110371000). Corrected blister exudate weights (without citrate buffer) were used as a proxy for blister volume. Exudates from the same time point were pooled and centrifuged at 400 × g for 5 min at RT, and the resulting pellet was resuspended in 1 mL of RPMI media (Cytiva, #SH30027.01) and kept on ice. 10 µL of the suspension was mixed with 0.4% Trypan Blue (1:1), and cells were counted using a Neubauer-improved counting chamber and light microscope. For RNA analysis, the cell suspension was centrifuged at 500 × g for 5 min at 4 °C. The pellet was resuspended, thoroughly vortexed and incubated in 1 mL of Qiazol (Qiagen, Cat#79306) for 15 min at RT, snap frozen in liquid nitrogen and stored at −80 °C. Supernatants were transferred to polypropylene tubes (BD, Cat#352063), centrifuged at 2000 × g for 10 min at RT, aliquoted, and stored at −80 °C.

#### Laser doppler imaging

Blood flow at the site of cantharidin application was measured at baseline (pre-application), 24 h and 72 h. Imaging was performed after blister harvest to avoid interference from exudate. The forearm was positioned 35 cm below the scanner and imaged using a laser Doppler imager (Moor LDI, Moor Instruments). Data were collected and visualized as a two-dimensional color-coded image, and processed using Moor Software V.6.2 (Moor Instruments), with the blister area manually defined as the region of interest (ROI). Blood flow was expressed as arbitrary perfusion units (PU), derived from Doppler frequency shifts reflecting red blood cell velocity and concentration (Motwani et al., 2016; Saez et al., 2005).

#### Inflammation-related proteomics in plasma and blister exudates

Following an overnight fast, venous blood was collected into K₂EDTA tubes and centrifuged at 2000 × g for 10 min (RT) at precisely 30 min post-collection. Plasma was aliquoted and snap frozen in liquid nitrogen and stored at −80°C until analysis. A total of 92 inflammation-related proteins were quantified in plasma and supernatants of blister exudates collected at 24 h and 72 h. Quantification was performed using the Target 96 Inflammation panel (v3026) via Olink proximity extension assay (Olink Proteomics, Uppsala, Sweden) at the SciLifeLab Clinical Biomarkers Facility (Uppsala, Sweden). Following the manufacturer’s protocol, 1 µl of sample was incubated with oligonucleotide-labeled antibody pairs. Target binding triggered DNA hybridization, extension, and high-throughput qPCR amplification using the BioMark HD system (Fluidigm Corporation). To eliminate inter-assay variability, all plasma and blister samples were processed on their own respective single plates. To account for sample-specific dilution, blister exudate concentrations were normalized using a correction factor derived from the total recovered volume relative to the initial 50 µL of citrate. Proteins with ≥ 30% values below limit of detection in all groups were excluded from downstream analyses.

#### RNA isolation

Total RNA was extracted from blister cells homogenized in Qiazol using the miRNeasy Mini Kit (Qiagen, Cat#217004) and treated with on-column DNase I per the manufacturer’s instructions. Venous blood was collected in PAXgene blood RNA tubes (Qiagen, PreAnalytiX, Cat#762165), incubated for 2 h at RT, and stored at −20 °C before transfer to −80 °C. RNA was isolated using PAXgene Blood miRNA Kit (Qiagen, Cat#763134) according to the manufacturer’s instructions. Briefly, samples were thawed on ice and centrifuged at RT for 10 min at 3000 × g. Pellets were washed twice with RNase-free water and centrifuged at RT for 10 min at 3000 × g. Samples were treated with Proteinase K (10 min, 55 °C, agitation 1000 rpm, Thermomixer Comfort, Eppendorf), followed by isopropanol-based RNA/DNA precipitation and DNase I digestion (15 min at RT). RNA was eluted in 40 µL BR5 buffer, with a repeated elution to increase total yield, then incubated for 5 min at 65 °C and chilled on ice. RNA concentration was measured on Nanodrop (ThermoFisher) and integrity was assessed using the RNA 6000 Pico Kit (Agilent, Cat#5067-1513) on Agilent 2100 Bioanalyzer (Agilent). Samples were stored at −80°C until analysis.

#### RNA Sequencing and Bioinformatic Analysis

Prior to library construction, the structural integrity of purified RNA was assessed. For bulk RNA sequencing, polyadenylated transcripts were reverse-transcribed into full-length cDNA using a template-switching protocol and barcoded primers containing Unique Molecular Identifiers (UMIs). Subsequent to cDNA amplification, libraries were prepared using an Illumina-compatible 10X Genomics construction kit. Following rigorous quality control, paired-end sequencing was executed on the Illumina NovaSeqX platform. The resulting raw FASTQ files were processed using STAR, with reads aligned to the human reference genome (GRCh38.108). While unique mapping was prioritized, unmapped reads were preserved for downstream alignment diagnostics. Gene-level quantification, including UMI extraction and deduplication, was performed via the STARsolo/GeneCounts module. Differential gene expression (DGE) was modeled in R (v4.5.2) utilizing the DESeq2 package (v1.50.2) (Love et al., 2014). To ensure robustness, we filtered the dataset to exclude low-abundance transcripts and restricted the analysis to protein-coding genes (annotated via biomaRt v2.66.0). To obtain more reliable effect size estimates and mitigate noise associated with low-count genes or high dispersion, log_2_fold change (LFC) values were stabilized using the ‘lfcshrink’ function with the ’apeglm’ adaptive shrinkage estimator. Significant differentially expressed genes (DEGs), defined by an absolute shrunken |LFC| > 0.5 and an adjusted p < 0.05, were subjected to functional enrichment via Metascape (v3.5.20260201) or clusterProfiler (v4.18.4) (Yu et al., 2012) targeting Gene Ontology Biological Process (GO:BP) terms. Shrunken LFC estimates were further utilized to generate four-way plots, enabling a comparative visualization of transcriptomic shifts across selected phenotypes. Visualizations were produced using EnhancedVolcano (v1.28.2), ComplexHeatmap (v2.26.0) (Gu et al., 2016), or GraphPad Prism (v10.6.1).

#### Flow cytometry

Flow cytometry was performed on whole blood and blister samples. Reagents were purchased from BD Biosciences unless stated otherwise. Briefly, red blood cell lysis was performed on 100 µL of EDTA blood using 3 mL ammonium chloride solution (80.2 g NH_4_CL, 8.4g NaHCO_3_ and 3.7 g EDTA in 1L ultrapure water, diluted 1:10 with deionized water) added in three 1 mL steps while vortexing. After a 15 min incubation at RT, samples were centrifuged at 320 × g, 20 °C for 5 min. Cells were washed once with 2 mL PBS. Next, 5 µL of BD Human FC block (2.5 µg/mL, Cat#564220) was added and incubated for 10 min at RT. Subsequently cells were stained using antibody cocktails (**Table S1**) prepared in Brilliant Stain Buffer Plus (Cat#566385). After a 15 min incubation in the dark, 1 mL of diluted BD FACS Lysing Solution (Cat#349202) was added and incubated for an additional 15 min in the dark. Cells were centrifuged at 320 × g, 20 °C for 5 min and washed twice with 2 mL PBS/HSA, then resuspended in PBS/HSA and stored at 4 °C in the dark until analysis. Blister cells were treated similarly, red blood cell lysis was omitted, and cells were centrifuged at 460 x g. Blister cells were additionally stained with a viability dye (FVS-450, Cat#562247) prior to FC block. Stainings were prepared in Brilliant Stain Buffer (Cat#563794). Events were acquired with the Penteon or BD FACSLyric using NovoExpress or FACSuite software. Subsequent analysis was performed using Flowjo v10.10. Gating strategies are shown in **Fig. S3**. To optimize the discrimination of CCR4- and CCR6-positive and -negative populations, the respective gates were initially established within the single-cell population and subsequently applied to the relevant T-cell subpopulations. Due to technical issues affecting a subset of samples, CD4^+^ T-cell subset data were not available for all participants.

#### Quantification and statistical analysis

A p value ≤ 0.05 was considered statistically significant. For high-dimensional transcriptomic and proteomic datasets, p values were adjusted for multiple testing using the Benjamini-Hochberg (BH) procedure to control the False Discovery Rate (FDR).

Baseline clinical characteristics and flow cytometry parameters were analyzed based on data type. Continuous variables were compared across groups using the non-parametric Kruskal-Wallis test, with pairwise differences evaluated via Dunn’s post-hoc test. Categorical variables were assessed using Fisher’s exact test. Comparisons for clinical parameters (**Table 1**) were further corrected for multiple comparisons using BH method. All statistical tests were two-tailed.

Plasma proteomic data were evaluated using one-way ANOVA followed by Tukey’s HSD post-hoc test. For blister exudate, protein abundance was analyzed using linear mixed-effects models (LMMs) to account for the longitudinal nature of the data and inter-individual variability. These models, adjusted for age and sex, incorporated metabolic phenotype and time point as fixed effects, with individuals included as random effects. Models were estimated using Restricted Maximum Likelihood (REML). To evaluate specific phenotypic differences, we calculated estimated marginal means (EMMs), and pairwise comparisons were performed for the metabolic phenotype × time point interaction. Degrees of freedom for inference were determined using the Kenward–Roger approximation. Selected comparisons were subjected to FDR correction and deemed significant at p ≤ 0.05. All analyses were implemented in the R environment (v4.5.2) utilizing the nlme (v3.1-166) and emmeans (v1.11.1) packages.

### Online supplemental material

Fig. S1 presents the flow chart of the study recruitment process. **Fig. S2** presents PLS-DA results and corresponding VIP analysis of blister exudates, along with supplemental RNAseq analyses. **Fig. S3** outlines the flow cytometry gating strategy for the different staining panels used in this study. **Fig. S4** presents PLS-DA results and corresponding VIP analysis of plasma samples. **Table S1** lists the antibodies used for flow cytometry in this study.

---

**Figure S1.**
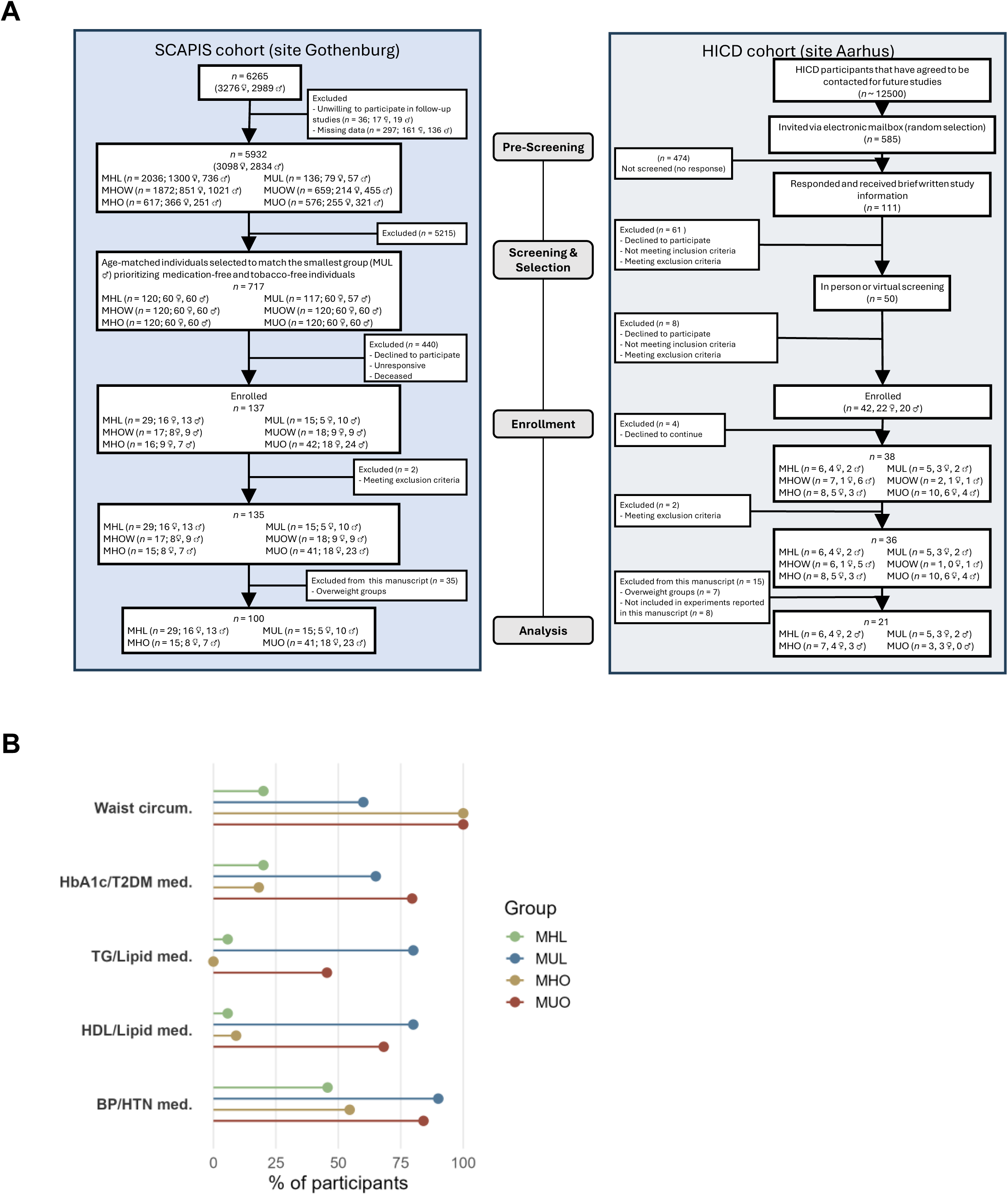
Recruitment flow and metabolic phenotype classification of study participants. **(A)** Recruitment flow of participants from the SCAPIS and HICD cohorts. Patient phenotyping into metabolically healthy lean, metabolically unhealthy lean, metabolically healthy obese, and metabolically unhealthy obese groups was based on predefined criteria described in the Materials and methods. In SCAPIS, participants were pre-stratified by metabolic phenotype. In 2020, all eligible metabolically unhealthy lean individuals and age-matched participants of both sexes from the remaining phenotype groups were selected for contact. Following enrollment, exclusions, and reassessment at the first study visit, participants were included in the final analytical cohort. In HICD, individuals who had consented to future contact were randomly invited, screened, and enrolled. After exclusion of individuals with overweight or incomplete experimental data, HICD participants were included in the analyses. **(B)** Proportion of participants fulfilling each of the five metabolic classification criteria within each phenotype group. Criteria include waist circumference, HbA1c and/or treatment for type 2 diabetes, triglycerides and/or lipid-lowering medication, HDL cholesterol and/or lipid-lowering medication, and blood pressure and/or antihypertensive treatment. Points indicate the percentage of participants meeting each criterion. Criteria and cutoffs are described in Methods. SCAPIS, Swedish CArdioPulmonary BioImage Study; HICD, Health in Central Denmark; MHL, metabolically healthy lean; MUL, metabolically unhealthy lean; MHO, metabolically healthy obese; MUO, metabolically unhealthy obese. ***Supplemental Figure S1 is related to*** ***Table 1***.

**Figure S2.**
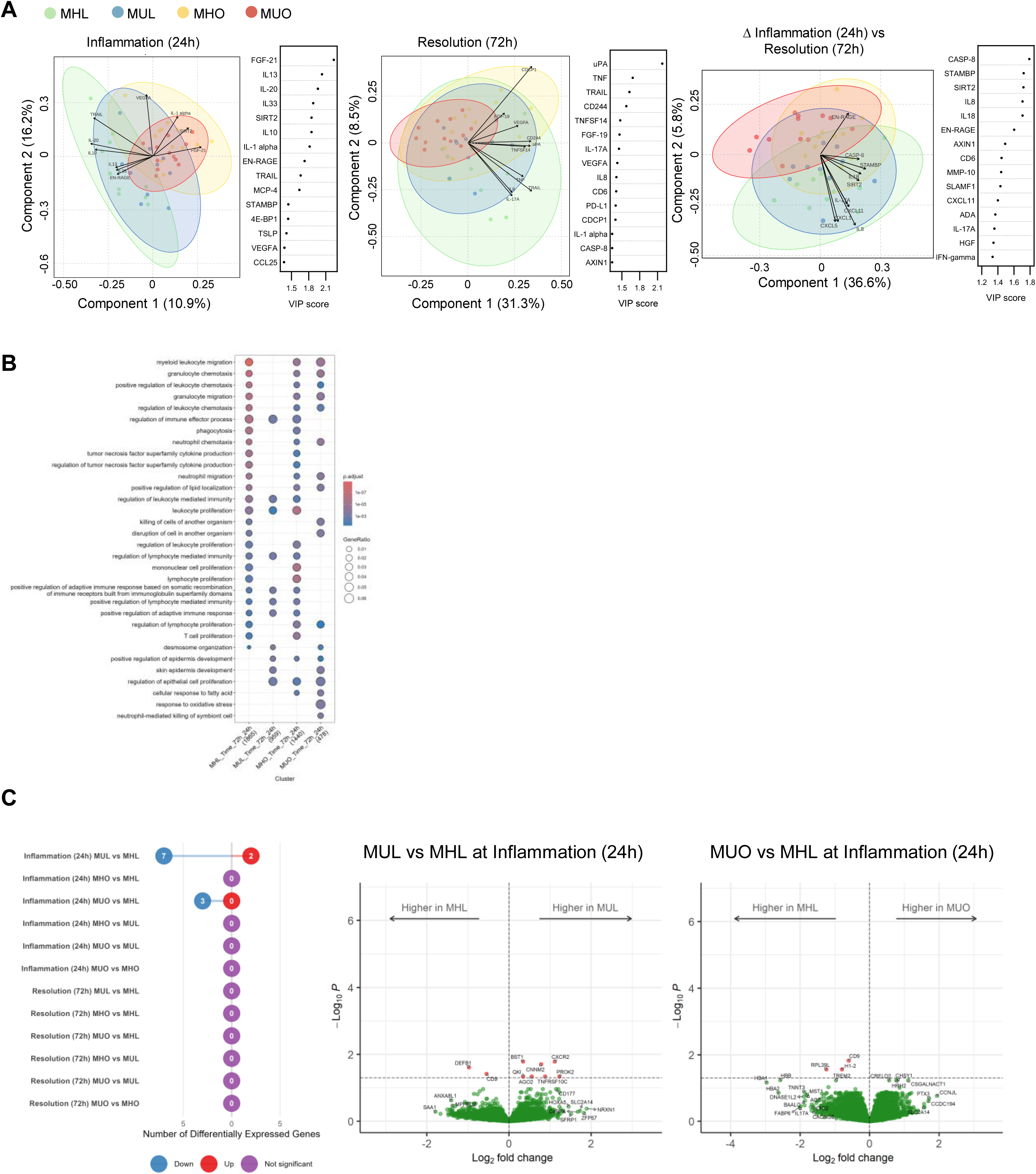
Blister exudate proteomics and blister-cell transcriptomics. **(A)** Partial Least Squares-Discriminant Analysis (PLS-DA) score plot of blister exudate protein expression levels measured using Olink Inflammation panel with key proteins driving group separation presented as Variable Importance in Projection (VIP) score, during the inflammation phase, the resolution phase, and the difference between the time points. **(B)** Gene Ontology Biological Process (GO:BP) terms identified by differential RNA-Seq gene expression analysis of blister 72h vs 24h time points (Fig. 1H) differing between metabolic phenotypes. **(C)** Left panel: Differential gene expression RNA-Seq analysis in peripheral tissue blister model per time point. Plot illustrating the number of regulated genes during the inflammation (24 h) and resolution (72 h) phase in direct comparisons across metabolic phenotypes. Right panel: Volcano plots illustrating global transcriptional changes in the metabolic phenotype and time point, where significant changes were identified. MUL compared to MHL (left panel) and MUO compared to MHL (right panel) during the inflammation phase (24h). The x-axis represents stabilized log2 fold changes (LFC) and the y-axis denotes log10 adjusted p values. Regulated genes are shown in red and are annotated. ***Supplemental Figure S2 is related to*** Figure 1.

**Figure S3.**
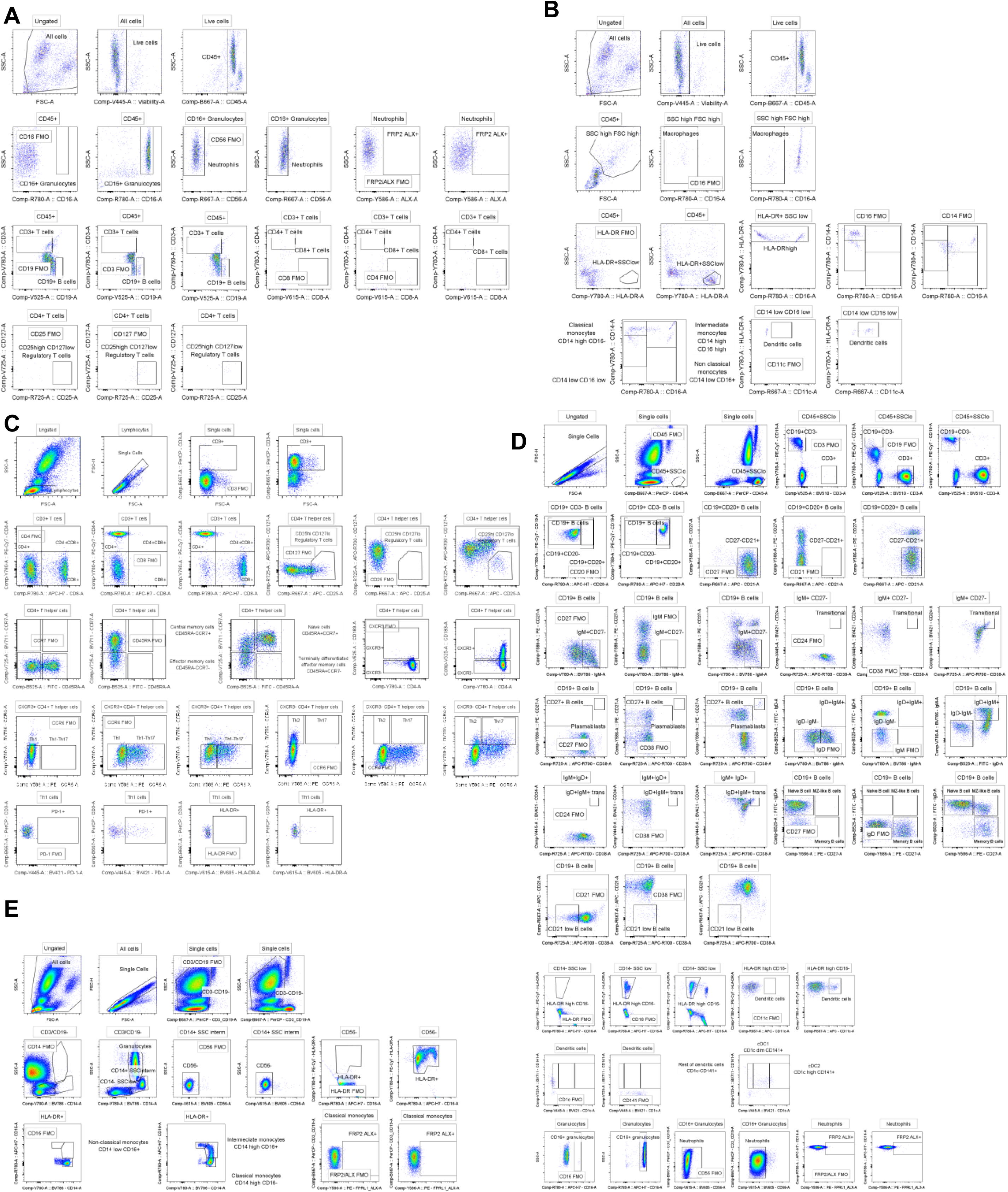
Gating strategy flow cytometry. **(A)** Gating strategy for identification of neutrophils, T-cells and B-cells in blister-derived leukocytes. **(B)** Gating strategy for identification of macrophages and monocytes in blister-derived leukocytes. **(C)** Gating strategy for T-cell populations in whole blood. Tc subsets were defined using the same gating approach as Th subsets. Expression of PD-1 and HLA-DR was assessed within each subset as indicated. **(D)** Gating strategy for B-cell populations in whole blood. CD27^+^, plasmablasts and transitional cells were gated within appropriate B cells subsets according to the same strategy as presented in the figure. **(E)** Gating strategy for identification of monocytes, dendritic cells and neutrophils in whole blood. Text boxes above each plot indicate the gating step used to generate the displayed population. Fluorescence minus one (FMO) controls and gate names are shown within each plot; where space was limited, gate names are indicated adjacent to the corresponding plot. ***Supplemental Figure S3 is related to*** ***Table 2-3*.**

**Figure S4.**
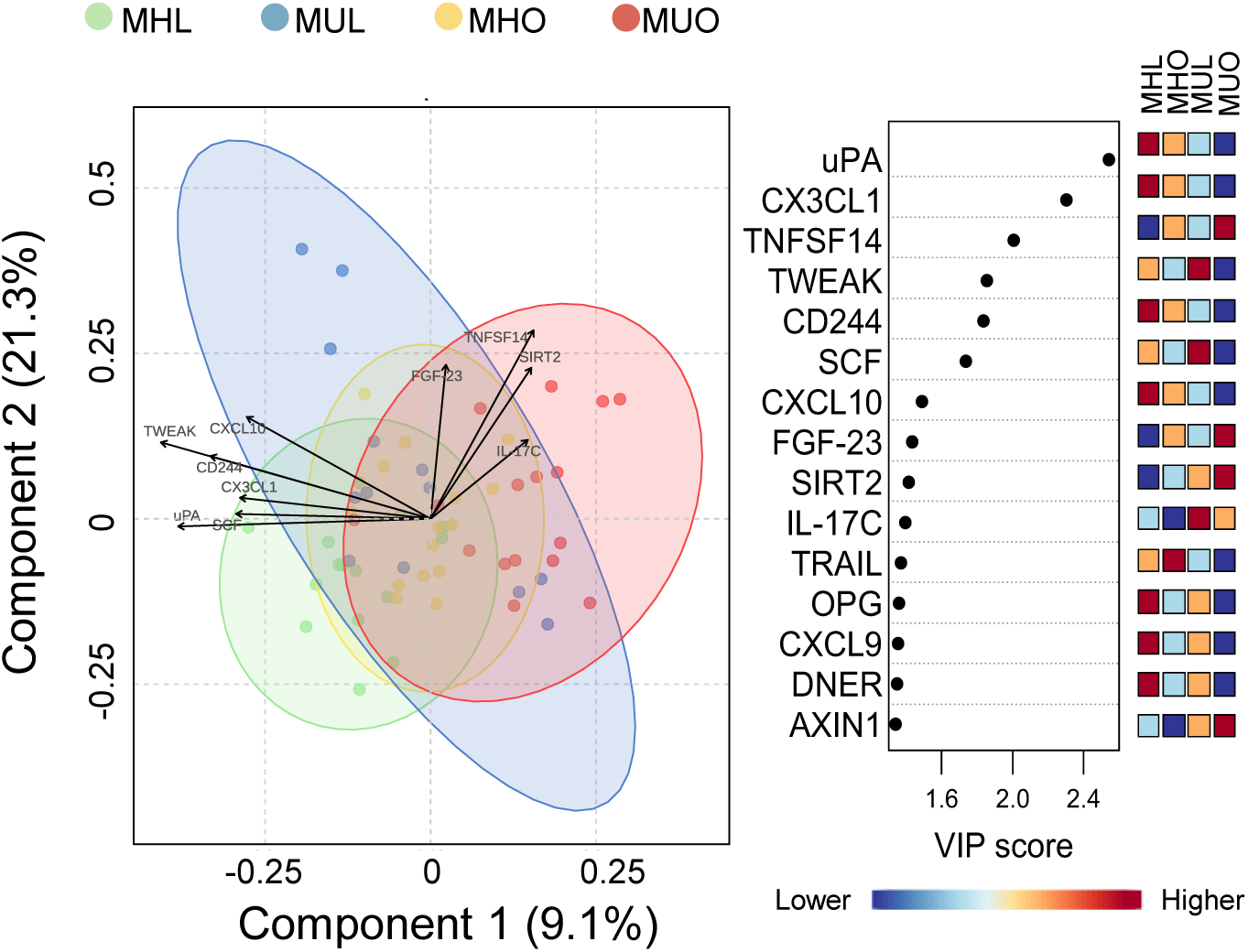
Partial least squares-discriminant analysis of plasma proteomics. Partial Least Squares-Discriminant Analysis (PLS-DA) score plot of plasma protein expression levels (NPX in log2 scale) related to inflammatory processes measured using the Olink Inflammation panel. Key proteins driving group differences are identified through Variable Importance in Projection (VIP) score analysis. ***Supplemental Figure S4 is related to*** Figure 2.

**Supplemental table 1.** Overview of antibodies used in the flow cytometry analyses

| Antigen | Dye | clone | Species | Host | Manufacturer | cat.no. |
| --- | --- | --- | --- | --- | --- | --- |
| CD11c | APC | B-ly6 | Human | Mouse | BD Biosciences | 559877 |
| CD127 | BV711 | HIL-7R-M21 | Human | Mouse | BD Biosciences | 563165 |
| CD14 | BV786 | M5E2 | Human | Mouse | BD Biosciences | 563698 |
| CD16 | APC-H7 | 3G8 | Human | Mouse | BD Biosciences | 560195 |
| CD19 | BV510 | SJ25C1 | Human | Mouse | BD Biosciences | 562947 |
| CD25 | APC-R700 | 2A3 | Human | Mouse | BD Biosciences | 565106 |
| CD3 | BV786 | SK7 | Human | Mouse | BD Biosciences | 563800 |
| CD56 (NCAM-1) | APC | B159 | Human | Mouse | BD Biosciences | 555518 |
| CD8 | BV605 | SK1 | Human | Mouse | BD Biosciences | 564116 |
| HLA-DR | PE-Cy7 | L243 | Human | Mouse | BD Biosciences | 335830 |
| CD127 | APC-R700 | HIL-7R-M21 | Human | Mouse | BD Biosciences | 565185 |
| CD183 (CXCR3) | BV510 | 1C6/CXCR3 | Human | Mouse | BD Biosciences | 740183 |
| CD194 (CCR4) | BV786 | 1G1 | Human | Mouse | BD Biosciences | 744141 |
| CD196 (CCR6) | PE | 11A9 | Human | Mouse | BD Biosciences | 559562 |
| CD197 (CCR7) | BV711 | 2-L1-A | Human | Mouse | BD Biosciences | 566752 |
| CD25 | APC | 2A3 | Human | Mouse | BD Biosciences | 340907 |
| CD279 (PD-1) | BV421 | EH12.1 | Human | Mouse | BD Biosciences | 562516 |
| CD45RA | FITC | HI100 | Human | Mouse | BD Biosciences | 555488 |
| CD8 | APC-H7 | SK1 | Human | Mouse | BD Biosciences | 641400 |
| HLA-DR | BV605 | G46-6 | Human | Mouse | BD Biosciences | 562845 |
| CD3 | PerCP | SK7 | Human | Mouse | BD Biosciences | 345766 |
| CD4 | PE-Cy7 | SK3 | Human | Mouse | BD Biosciences | 348809 |
| CD19 | PE-Cy7 | SJ25C1 | Human | Mouse | BD Biosciences | 341113 |
| CD20 | APC-H7 | L27 | Human | Mouse | BD Biosciences | 641414 |
| CD21 | APC | B-ly4 | Human | Mouse | BD Biosciences | 559867 |
| CD24 | BV421 | ML5 | Human | Mouse | BD Biosciences | 562789 |
| CD27 | PE | L128 | Human | Mouse | BD Biosciences | 340425 |
| CD3 | BV510 | SK7 | Human | Mouse | BD Biosciences | 740202 |
| CD38 | APC-R700 | HIT2 | Human | Mouse | BD Biosciences | 564979 |
| IgD | FITC | IA6-2 | Human | Mouse | BD Biosciences | 555778 |
| IgM | BV786 | G20-127 | Human | Mouse | BD Biosciences | 740998 |
| CD45 | PerCP | 2D1 | Human | Mouse | BD Biosciences | 345809 |
| CD141 | BV711 | 1A4 | Human | Mouse | BD Biosciences | 563155 |
| CD16 | APC-H7 | 3G8 | Human | Mouse | BD Biosciences | 560195 |
| CD19 | PerCP | 4G7 | Human | Mouse | BD Biosciences | 345790 |
| CD1c | BV421 | F10/21A3 | Human | Mouse | BD Biosciences | 565050 |
| CD56 (NCAM-1) | BV605 | B159 | Human | Mouse | BD Biosciences | 740405 |
| CD56 (NCAM-1) | BV605 | NCAM16.2 | Human | Mouse | BD Biosciences | 562780 |
| HLA-DR | PE-Cy7 | L243 | Human | Mouse | BD Biosciences | 335830 |
| CD14 | BV786 | M5E2 | Human | Mouse | BD Biosciences | 563698 |
| FPRL1-FPRL2 | PE | 304405 | Human | Mouse | RD System | FAB3479P |

